# Identify potent SARS-CoV-2 main protease inhibitors via accelerated free energy perturbation-based virtual screening of existing drugs

**DOI:** 10.1101/2020.03.23.004580

**Authors:** Zhe Li, Xin Li, Yi-You Huang, Yaoxing Wu, Runduo Liu, Lingli Zhou, Yuxi Lin, Deyan Wu, Lei Zhang, Hao Liu, Ximing Xu, Kunqian Yu, Yuxia Zhang, Jun Cui, Chang-Guo Zhan, Xin Wang, Hai-Bin Luo

## Abstract

Coronavirus disease 2019 (COVID-19) pandemic caused by severe acute respiratory syndrome coronavirus 2 (SARS-CoV-2) has become a global crisis. There is no therapeutic treatment specific for COVID-19. It is highly desirable to identify potential antiviral agents against SARS-CoV-2 from existing drugs available for other diseases and, thus, repurpose them for treatment of COVID-19. In general, a drug repurposing effort for treatment of a new disease, such as COVID-19, usually starts from a virtual screening of existing drugs, followed by experimental validation, but the actual hit rate is generally rather low with traditional computational methods. Here we report a new virtual screening approach with accelerated free energy perturbation-based absolute binding free energy (FEP-ABFE) predictions and its use in identifying drugs targeting SARS-CoV-2 main protease (M^pro^). The accurate FEP-ABFE predictions were based on the use of a new restraint energy distribution (RED) function designed to accelerate the FEP-ABFE calculations and make the practical FEP-ABFE-based virtual screening of the existing drug library possible for the first time. As a result, out of twenty-five drugs predicted, fifteen were confirmed as potent inhibitors of SARS-CoV-2 M^pro^. The most potent one is dipyridamole (K_i_=0.04 μM) which has showed promising therapeutic effects in subsequently conducted clinical studies for treatment of patients with COVID-19. Additionally, hydroxychloroquine (K_i_=0.36 μM) and chloroquine (K_i_=0.56 μM) were also found to potently inhibit SARS-CoV-2 M^pro^ for the first time. We anticipate that the FEP-ABFE prediction-based virtual screening approach will be useful in many other drug repurposing or discovery efforts.

**Significance Statement:** Drug repurposing effort for treatment of a new disease, such as COVID-19, usually starts from a virtual screening of existing drugs, followed by experimental validation, but the actual hit rate is generally rather low with traditional computational methods. It has been demonstrated that a new virtual screening approach with accelerated free energy perturbation-based absolute binding free energy (FEP-ABFE) predictions can reach an unprecedently high hit rate, leading to successful identification of 16 potent inhibitors of SARS-CoV-2 main protease (M^pro^) from computationally selected 25 drugs under a threshold of K_i_ = 4 μM. The outcomes of this study are valuable for not only drug repurposing to treat COVID-19, but also demonstrating the promising potential of the FEP-ABFE prediction-based virtual screening approach.

---

The ongoing pandemic of coronavirus disease 2019 (COVID-19)(1, 2) caused by severe acute respiratory syndrome coronavirus 2 (SARS-CoV-2, also known as 2019-nCoV), has become a global crisis. To date, there is no specific treatment or vaccine for COVID-19. Thus, there is an urgent need to repurpose drugs for treatment of COVID-19.(3) The SARS-CoV-2 replicase gene (Orf1) encodes two overlapping translation products, polyproteins 1a and 1ab (pp1a and pp1ab), which mediate all of the functions required for the viral replication. The main protease (M^pro^) as a key enzyme for the viral replication is initially released by the autocleavage of pp1a and pp1ab. Then, M^pro^ cleaves pp1a and pp1ab to release the functional proteins nsp4-nsp16 that are necessary for the viral replication.(4) In view of the essential functions of M^pro^ in the viral life cycle and its high level of conservation, SARS-CoV-2 M^pro^ is a naturally attractive target for treatment of COVID-19. Hence, there have been efforts to identify therapeutic candidates targeting M^pro^ using various virtual screening methods based on pharmacophore, molecule docking, and molecular simulations.(5) As a result of the reported efforts, six drugs were found to inhibit SARS-CoV-2 M^pro^ with IC_50_ ranging from 0.67 to 21.4 μM.^5^ There have been also drug repurposing efforts associated with other potential targets of SARS-CoV-2.^5-14^

In general, a drug repurposing effort for treatment of a new disease, such as COVID-19, usually starts from a virtual screening of existing drugs through computational modeling and simulations, followed by experimental validation. However, the actual hit rate of a virtual screening using traditional computational methods has been rather low, with vast majority of computationally predicted drug candidates being false positives, because it is difficult to reliably predict protein-ligand binding free energies. Most recently, Gorgulla *et al*.(6) reported an interesting new virtual screening platform, called VirtualFlow, used to screen numerous compounds in order to identify inhibitors of Kelch-like ECH-associated protein 1 (KEAP1), but the hit rate was still not very high. Within 590 compounds predicted by the virtual screening, 69 were found to be KEAP1 binders (with a hit rate of ~11.7% for detectable binding affinity), and 10 of these compounds were confirmed to be displacers of nuclear factor erythroid-derived 2-related factor 2 (NRF2) with a half-maximum inhibitory concentration (IC_50_) < 60 μM (with a hit rate of ~1.4% under the threshold of IC_50_ < 60 μM).(6) Obviously, the hit rate of a virtual screening is dependent on the reliability and accuracy of the receptor-ligand binding free energy predictions used in the virtual screening process. So, the key to the success of a virtual screening effort is use of a reliable computational approach to accurately predict binding free energies.

The free energy perturbation (FEP) simulation of intermolecular interactions (7, 8) is recognized a reliable method for binding free energy calculations with satisfactory accuracy,(7–18) but the traditional FEP method was limited to simulating some minor structural changes of ligands for the relative binding free energy (RBFE) calculations.(9, 19) The RBFE calculations can be used to guide lead optimization starting from a promising lead compound (or hit),(9, 19–22) but not suitable for virtual screening of completely different molecular structures to identify new hits for drug repurposing. For the virtual screening to identify new hits or leads, it is necessary to predict absolute binding free energy (ABFE) for each ligand binding with the target without the requirement to use any reference ligand structure. The FEP-ABFE approach has the advantage of predicting binding affinities between ligands and their targets more accurately than conventional computational methods, such as pharmacophore, molecule docking, and molecular simulations.(23) However, the previously used FEP-ABFE calculations are extremely expensive and time-consuming and, thus, not suitable for virtual screening purposes (that required to screen a large number of compounds).(24, 25)

To make the FEP-ABFE approach practically feasible for our virtual screening and drug repurposing effort, here we report a new algorithm using a restraint energy distribution (RED) function to accelerate the FEP-ABFE prediction and its first application to a drug repurposing effort which targets SARS-CoV-2 M^pro^. Our FEP-ABFE prediction-based virtual screening (which predicted 25 drugs as potential inhibitors of SARS-CoV-2 M^pro^) was followed by *in vitro* activity assays, confirming that 15 out of the 25 drugs can potently inhibit SARS-CoV-2 M^pro^ with 0.04 to 3.3 μM (with a remarkably high hit rate of 60% under a threshold of K_i_ = 4 μM); nine drugs have K_i_ < 1 μM (with a submicromolar hit rate of 36%). Particularly, among these drugs, the most potent inhibitor of SARS-CoV-2 M^pro^ is dipyridamole (DIP, K_i_ = 0.04 μM). Following the computational prediction and *in vitro* activity validation, DIP was tested for its antiviral activity against SARS-CoV-2 *in vitro* and in clinical studies for treatment of patients with COVID-19, and the preliminary clinical data are promising for its actual therapeutic effects. While the clinical data are reported separately elsewhere(26) to timely guide further clinical studies and possibly practical clinical application, we describe and discuss in this report the detailed computational and *in vitro* activity results of DIP along with other promising drugs identified. The encouraging outcomes suggest that the FEP-ABFE prediction-based virtual screening is a truly promising approach to drug repurposing.

## Results and Discussion

### Identification of potent SARS-CoV-2 M^pro^ inhibitors for drug repurposing

Prior to the virtual screening for drug repurposing, the accuracy of the accelerated FEP-ABFE prediction protocol was validated by using three different protein targets (BRD4, HIV-1 protease, and human factor Xa) and 28 ligands with diverse chemical scaffolds. According to the validation data, given in Supporting Information (SI) section S7, the accelerated FEP-ABFE algorithm can achieve a high accuracy for the ABFE predictions. So, in order to identify potent SARS-CoV-2 M^pro^ inhibitors, we first carried out the FEP-ABFE based virtual screening of all existing drugs, followed by *in vitro* activity assays, as shown in Figure 1.

**Figure 1.**
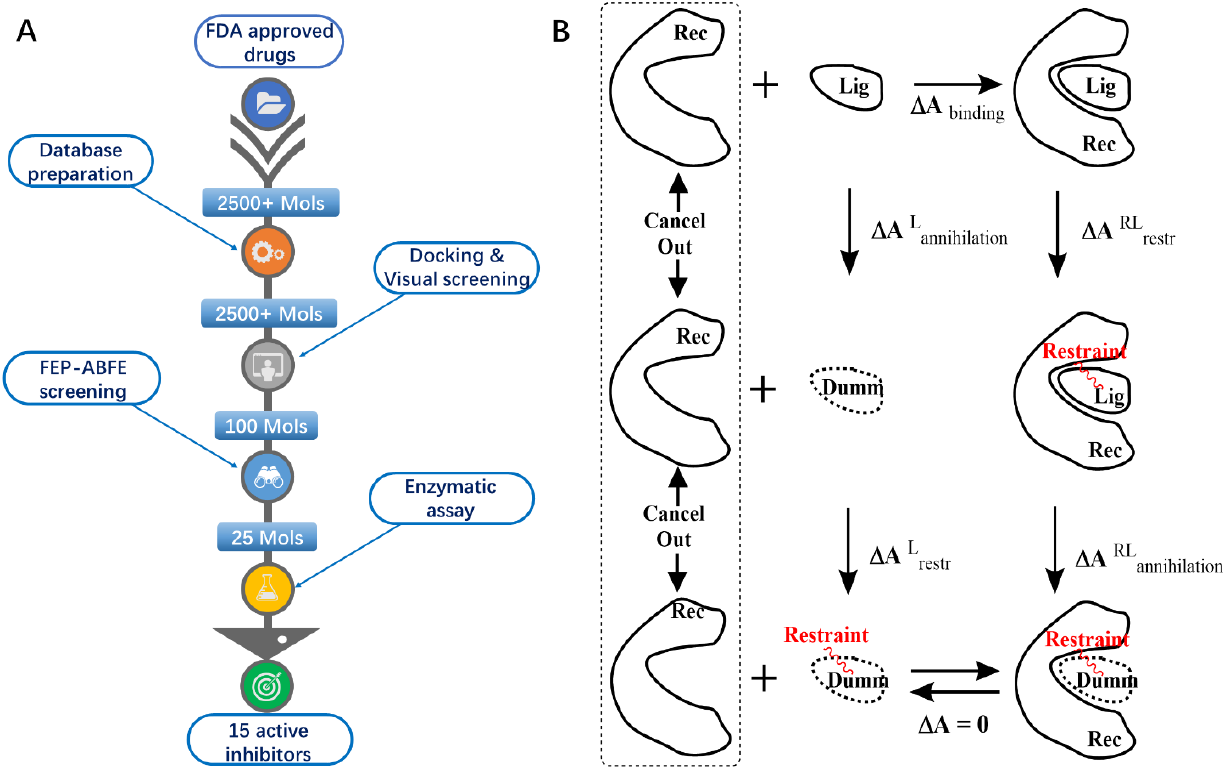
The FEP-ABFE based screening for the drug repurposing targeting SARS-CoV-2 M^pro^. (A) The schedule of FEP-ABFE based screening. (B) Thermodynamic cycle used for the FEP-ABFE calculations.

Specifically, after all the existing drugs were docked into the binding site of SARS-CoV-2 M^pro^, 100 molecules that had specific interactions with the six key amino acid residues, Cys145, His41, Ser144, His163, Gly143, and Gln166, were subjected to further FEP-ABFE calculations. Among these 100 drugs, 49, 46, and 5 were electrically neutral, negatively charged, and positively charged, respectively. Since the FEP method is known to encounter systematic errors when the ligands are not electrically neutral, the drugs selected on the basis of the FEP-ABFE results were grouped by their formal charges to ensure that the error is cancelled within each group. In each group, the top 20% to 40% of the molecules were selected based on their ABFE values. As a result, 25 drugs were selected for subsequent *in vitro* experimental activity testing. According to the *in vitro* results, 15 out of these 25 drugs exhibited considerable potency of inhibiting SARS-CoV-2 M^pro^ (Figures 2 and S8). DIP was found to be the most potent inhibitor, with K_i_ = 0.04 μM. Following the computational prediction and *in vitro* activity confirmation, DIP was further tested for its antiviral activity against SARS-CoV-2, demonstrating that DIP dose-dependently suppressed the SARS-CoV-2 replication with EC_50_ = 0.1 μM. The antiviral activity was consistent with the inhibitory activity against M^pro^. In addition, DIP was also tested clinically in treatment of patients with COVID-19, resulting in promising therapeutic data that are reported separately elsewhere (along with the raw antiviral activity data)(26) due to the urgent need of further clinical studies and possibly practical clinical application.

**Figure 2.**
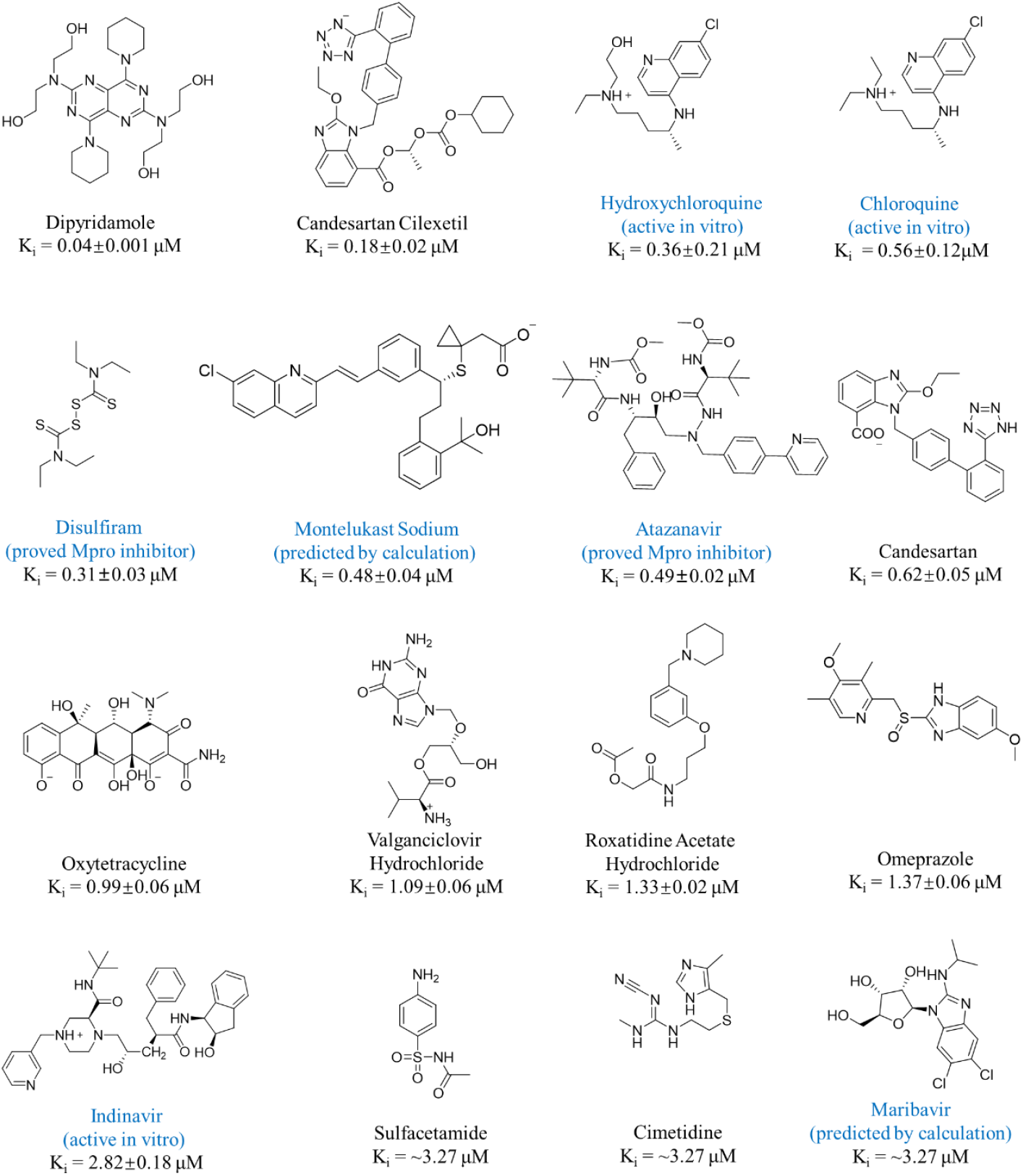
Molecular structures and K_i_ values of 16 confirmed SARS-CoV-2 M^pro^ inhibitors. The seven compounds in blue were also proposed as potential treatments for patients with COVID. Within the seven compounds, disulfiram and atazanavir were reported to be SARS-CoV-2 M^pro^ inhibitors with the reported IC_50_ listed in Table 1;(31, 32) hydroxychloroquine, chloroquine, and indinavir were reported to be active *in vitro* against COVID-19, but their molecular targets were not reported;(27–29) montelukast sodium and maribavir was only predicted by calculations(29, 30) without experimental activity data reported. Disulfiram served as the positive control for the *in vitro* activity (its IC_50_ value is 5.72 μM in the literature and 4.7 μM in this work when the concentration of the same substrate used was as high as 20 μM).

The FEP-ABFE results calculated for all the confirmed potent SARS-CoV-2 M^pro^ inhibitors are given in Table 1 in comparison with the subsequently determined experimental activity data. As seen in Table 1, 13 out of the 15 FEP-ABFE predicted binding free energies were within 2 kcal/mol of the corresponding experimental values, and for the other two (disulfiram and maribavir), the deviations were all about 2.2 kcal/mol. Specially for disulfiram, according to its molecular structure, it might be a covalent inhibitor of M^pro^, which could be part of the reason of the relatively larger computational error. However, further studies are needed for disulfiram to draw a more reliable conclusion. Overall, for the 15 protein-ligand binding complexes, the mean unsigned error (MUE) was about 1.2 kcal/mol. For comparison, we also carried out the MM-PBSA and MM-GBSA calculations on the 15 binding complexes, and the MUE values for both of the two methods were larger than 17.0 kcal/mol. Thus, the FEP-ABFE method is indeed much more accurate than both the MM-PBSA and MM-GBSA methods for the drug repurposing prediction.

**Table 1.** Summary of the FEP-ABFE, MM-PBSA, and MM-GBSA calculation results (in kcal/mol) for the experimentally confirmed SARS-CoV-2 M^pro^ inhibitors. The unsigned error (UE) and mean unsigned error (MUE) values are also given. ΔG_exp_ values were calculated from their corresponding K_i_ values.

| Name | IC<sub>50</sub> (μM)<sup>d</sup> | K<sub>i</sub> (μM)<sup>b</sup> | $\Delta G_{\text{exp}}$ | $\Delta G_{\text{FEP-ABFE}}$ | UE<sub>FEP-ABFE</sub><sup>c</sup> | $\Delta G_{\text{MM-PBSA}}$ | UE<sub>MM-PBSA</sub><sup>c</sup> | $\Delta G_{\text{MM-GBSA}}$ | UE<sub>MM-GBSA</sub><sup>c</sup> |
| --- | --- | --- | --- | --- | --- | --- | --- | --- | --- |
| Dipyridamole (DIP) | 0.6±0.01 | 0.04±0.001 | -10.1 | -8.6 | 1.5 | -23.7 | 13.6 | -34.5 | 24.4 |
| Candesartan cilexetil | 2.8±0.3 | 0.18±0.02 | -9.2 | -8.4 | 0.8 | -36.1 | 26.9 | -38.3 | 29.1 |
| Hydroxychloroquine | 2.9±0.3 | 0.36±0.21<sup>a</sup> | -8.7 | -9.8 | 0.7 | -24.0 | 14.9 | -28.7 | 19.6 |
| Chloroquine | 3.9±0.2 | 0.56±0.12<sup>a</sup> | -8.5 | -10.0 | 1.5 | -24.6 | 16.1 | -33.2 | 24.7 |
| Disulfiram | 4.7±0.4 (9.35±0.18)<sup>e</sup> | 0.31±0.03 | -8.8 | -6.6 | 2.2 | -23.0 | 14.2 | -24.3 | 15.5 |
| Montelukast sodium | 7.3±0.5 | 0.48±0.04 | -8.6 | -7.5 | 1.1 | -39.5 | 30.9 | -41.5 | 32.9 |
| Atazanavir | 7.5±0.3 (10)<sup>e</sup> | 0.49±0.02 | -8.6 | -7.6 | 1.0 | -33.0 | 24.4 | -39.2 | 30.6 |
| Oxytetracycline | 15.2±0.9 | 0.99±0.06 | -8.2 | -8.9 | 0.7 | -10.0 | 1.8 | -14.6 | 6.4 |
| Valacyclovir hydrochloride | 16.7±0.9 | 1.09±0.06 | -8.1 | -6.2 | 1.9 | -20.8 | 12.7 | -18.3 | 10.2 |
| Roxatidine acetate hydrochloride | 20.3±0.4 | 1.33±0.02 | -8.0 | -7.1 | 0.9 | -29.2 | 21.2 | -30.5 | 22.5 |
| Omeprazole | 21.0±1.0 | 1.37±0.06 | -8.0 | -6.5 | 1.5 | -22.3 | 14.3 | -24.4 | 16.4 |
| Indinavir | 43.1±2.8 | 2.82±0.18 | -7.6 | -8.3 | 0.7 | -28.8 | 21.2 | -35.4 | 27.8 |
| Sulfacetamide | ~50 | ~3.27 | -7.5 | -7.0 | 0.5 | -14.8 | 7.3 | -13.9 | 6.4 |
| Cimetidine | ~50 | ~3.27 | -7.5 | -6.0 | 1.5 | -26.2 | 18.7 | -27.8 | 20.3 |
| Maribavir | ~50 | ~3.27 | -7.5 | -5.3 | 2.2 | -25.4 | 19.5 | -31.0 | 25.7 |
| MUE | - | - | - | - | 1.2 | - | 17.2 | - | 20.8 |

<sup>a</sup> $K_i$ values for hydroxychloroquine and chloroquine were determined using the Dixon plots using the data in Figure 3.
<sup>b</sup> $K_i$ values for other molecules were converted from IC<sub>50</sub> based on the assumption of the competitive inhibition without covalent binding.
<sup>c</sup> UE<sub>FEP-ABFE</sub> = $|\Delta G_{\text{FEP-ABFE}} - \Delta G_{\text{exp}}|$ ; UE<sub>MM-PBSA</sub> = $|\Delta G_{\text{MM-PBSA}} - \Delta G_{\text{exp}}|$ ; UE<sub>MM-GBSA</sub> = $|\Delta G_{\text{MM-GBSA}} - \Delta G_{\text{exp}}|$
<sup>d</sup> IC<sub>50</sub> values when the substrate concentration was 20 μM.
<sup>e</sup> IC<sub>50</sub> values in the brackets are obtained from other published works, and the published values are close to our experiment results.(27-29)

Notably, candesartan cilexetil with K_i_ = 0.18 μM against SARS-CoV-2 M^pro^ is a prodrug for its labeled use (treatment of hypertension and congestive heart failure). Hence, we also computationally and experimentally examined its metabolite, candesartan (the active drug corresponding to the prodrug for the labeled use) which was not in the drug library screened. Interestingly, candesartan was also predicted and confirmed as a potent inhibitor of SARS-CoV-2 M^pro^, with a slightly lower inhibitory activity against SARS-CoV-2 M^pro^ (Ki = 0.62 μM). So, it is interesting to note that for potential treatment of patients with COVID-19, the prodrug candesartan cilexetil would serve as a more active molecular species against SARS-CoV-2 M^pro^ compared to candesartan itself.

Altogether, a total of 16 potent inhibitors of SARS-CoV-2 M^pro^ were identified in this study, and their molecular structures and *in vitro* inhibitory activity data are shown in Figures 2 and S8. Among these 16 compounds, nine (with names shown in black in Figure 2) were identified as potential candidate treatments of patients with COVID-19 for the first time in this study. The remaining seven drugs, including hydroxychloroquine, chloroquine, disulfiram, montelukast sodium, atazanavir, indinavir, and maribavir, were also proposed as potential candidate treatments for patients with COVID-19 in previous studies.(27, 29–32) However, within these seven drugs, only disulfiram and atazanavir were previously identified as SARS-CoV-2 M^pro^ inhibitors, whereas the other five drugs were either reported to be active *in vitro* against SARS-CoV-2 without knowing the specific targets or predicted by computational modeling only without knowing their actual experimental activity. All these drugs were confirmed to be potent SARS-CoV-2 M^pro^ inhibitors in this study. Overall, a total of 14 compounds were confirmed as potent SARS-CoV-2 M^pro^ inhibitors for the first time in this study.

Within the SARS-CoV-2 M^pro^ inhibitors identified, DIP is the most potent one with K_i_ = 0.04 μM (or 40 nM). The computationally modeled structure of DIP binding with SARS-CoV-2 is depicted in Figure S9 (showing the roles of key residues of the protease, including Thr25, Asn142, Gly143, Ser144, His163, and Glu166, for binding with DIP).

### Molecular mechanism for the antiviral activity of chloroquine and hydroxychloroquine against SARS-CoV-2

Notably, chloroquine and hydroxychloroquine are currently under clinical trials for treatment of patients with COVID-19. Particularly, chloroquine was reported to inhibit SARS-CoV-2 with EC_50_ of 0.1 ~ 1.13 μM,(26)(27) although the exact molecular mechanism and drug target(s) have not been confirmed. Concerning the molecular mechanism for their known antiviral activity, chloroquine or hydroxychloroquine was previously proposed to inhibit acidification of endosome and viral endocytosis.(33, 34) However, vesicular stomatitis virus (VSV), which was a model virus belonging to *Rhabdoviridae* and had a similar endocytosis process as coronavirus, was not as sensitive as SARS-CoV-2 to hydroxychloroquine and chloroquine (Figure S10); no significant inhibition was observed for hydroxychloroquine or chloroquine at a concentration 6.25 μM. Comparing to VSV, coronavirus is much more sensitive to chloroquine and hydroxychloroquine. Hydroxychloroquine inhibited SARS-CoV-2 at EC_50_ of 0.72 μM and chloroquine reduced SARS-CoV replication to 53% at 1.0 μM.(35) We were always wondering if chloroquine and its analogue hydroxychloroquine would directly target a viral protein of coronavirus. Here, we demonstrated in this report for the first time that chloroquine and its analogues inhibited the main protease (M^pro^) activity, which is an essential and conserved enzyme in *Coronaviridae*. Chloroquine and hydroxychloroquine are potent inhibitors of SARS-CoV-2 M^pro^ with K_i_ = 0.56 and 0.36 μM, respectively (see Figure 3). Here, we cautiously concluded that chloroquine and hydroxychloroquine prevented SARS-CoV-2 infection by inhibition of M^pro^ in addiction to the well-known mechanism of abrogation of viral endocytosis. Moreover, norovirus, which belonged to *Caliciviridae* and encoded a viral 3C-like protein similar to M^pro^ of coronavirus, was hypersensitive to chloroquine treatment.(36) It furtherly supported that chloroquine and its analogues would inhibit viral 3C-like protease and inhibit viral replication. The K_i_ value of 0.36 μM for hydroxychloroquine against SARS-CoV-2 M^pro^ is slightly lower than the reported EC_50_ of 0.72 μM against SARS-CoV-2, which is consistent with the possible molecular mechanism that the antiviral activity of hydroxychloroquine against SARS-CoV-2 is mainly due to the inhibitory activity against SARS-CoV-2 M^pro^. Overall, hydroxychloroquine or chloroquine is expected to have both some beneficial effect associated with its antiviral activity due to the SARS-CoV-2 M^pro^ inhibition and adverse side effects associated with other complicated mechanisms of the drug.

Further, in light of our finding that these drugs are potent SARS-CoV-2 M^pro^ inhibitors, it would be interesting to design hydroxychloroquine analogs that can more potently and selectively inhibit SARS-CoV-2 M^pro^ without the unwanted adverse effects of hydroxychloroquine. Similar drug development strategies may also apply to development of analogs of other confirmed SARS-CoV-2 M^pro^ inhibitors such as DIP and candesartan cilexetil with further improved potency and selectivity for SARS-CoV-2 M^pro^.

**Figure 3.**
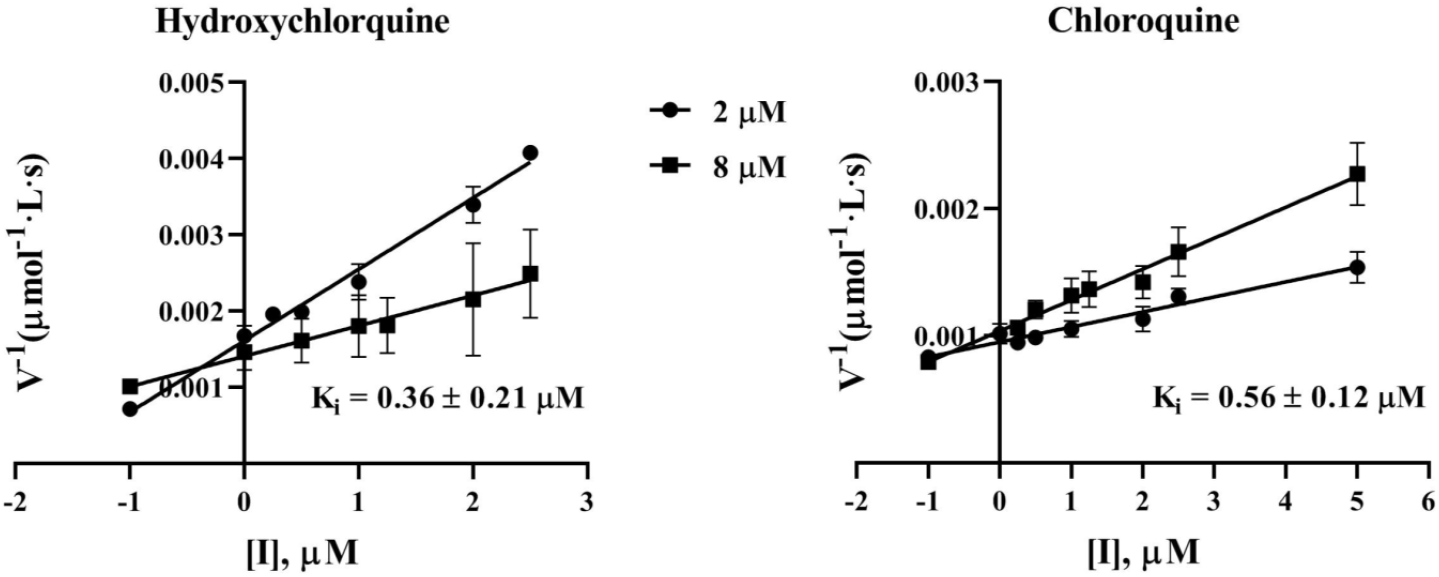
Chloroquine and hydroxychloroquine were identified as SARS-CoV-2 M^pro^ inhibitors with K_i_ = 0.56 and 0.36 μM, respectively. K_i_ was determined according to the enzymatic Kinetics using the Dixon plots.

## Conclusion

By using the accelerated free energy perturbation-based absolute binding free energy (FEP-ABFE) predictions for drug repurposing targeting SARS-CoV-2 M^pro^, followed by experimental validation, we successfully identified a total of 16 potent inhibitors of SARS-CoV-2 M^pro^ from existing drugs, including 14 SARS-CoV-2 M^pro^ inhibitors that were confirmed (with K_i_ = 0.04 to 3.3 μM) for the first time in this study. The identified most potent SARS-CoV-2 M^pro^ inhibitor is dipyridamole (with K_i_ = 0.04 μM) which is currently under clinical studies for treatment of patients with COVID-19, with the promising therapeutic effects reported in a separate report. Among other newly identified SARS-CoV-2 M^pro^ inhibitors, prodrug candesartan cilexetil and the corresponding drug candesartan both can potently inhibit SARS-CoV-2 M^pro^. Interestingly, prodrug candesartan cilexetil (with K_i_ = 0.18 μM) is even more potent than candesartan itself (with K_i_ = 0. 62 μM) for inhibiting SARS-CoV-2 M^pro^.

Additionally, hydroxychloroquine (Ki = 0.36 μM) and chloroquine (Ki = 0.56 μM) were found to potently inhibit SARS-CoV-2 M^pro^ for the first time in this study, suggesting that the previously known antiviral activity of hydroxychloroquine or chloroquine might be mainly due to the inhibitory activity against SARS-CoV-2 M^pro^, in addition to other well-known mechanisms. Further, based on the finding that these drugs are potent SARS-CoV-2 M^pro^ inhibitors, it would be interesting to design hydroxychloroquine analogs that can more potently and selectively inhibit SARS-CoV-2 M^pro^ to improve its antiviral activity and avoid the unwanted adverse effects of hydroxychloroquine associated with other mechanisms. Similarly, the identified other drugs, such as dipyridamole and candesartan cilexetil *etc*., can also be used as promising starting drug structures to design new drug candidates with further improved potency and selectivity for SARS-CoV-2 M^pro^.

In summary, the virtual screening through accelerated FEP-ABFE predictions has demonstrated an excellent accuracy, with a remarkably high hit rate of 60% under a threshold of K_i_ = 4 μM. We anticipate that the FEP-ABFE prediction-based virtual screening approach will be useful in many other drug repurposing or discovery efforts.

## Methods

### Virtual screening based on accelerated FEP-ABFE approach

The accelerated FEP-ABFE approach was based on the use of a new restraint energy distribution (RED) function. The RED function was derived to accelerate the FEP-ABFE calculations, and the accelerated FEP-ABFE approach are extensively tested and evaluated; see details given in Supporting Information (SI) sections S1 to S7. Briefly, the RED function accelerated FEP-ABFE approach was extensively tested and showed remarkable accuracy. Compared to the previously reported FEP-ABFE approaches which normally use 42 λ values(24, 25), the RED function accelerated FEP-ABFE can be calculated by using just 16 λ values. With such acceleration, the application of FEP-ABFE calculation in virtual screening was made possible. The accuracy of the accelerated 16-λ-FEP-ABFE calculation was then tested against 28 ligands with diverse chemical scaffolds, as given in SI section S7. The test results suggested that the accelerated FEP-ABFE algorithm can achieve a remarkable accuracy, which encouraged us to perform the FEP-ABFE prediction-based practical virtual screening to identify SARS-CoV-2 M^pro^ inhibitors for drug repurposing.

During the virtual screening, molecular docking was first performed by using the crystal structure (PDB ID: 6LU7)(31) of SARS-CoV-2 M^pro^ which causes COVID-19. More than 2500 small molecules in the existing drug library (including all FDA-approved drugs) were screened by docking method, and 100 ligands were selected by molecular docking and further evaluated by RED function accelerated FEP-ABFE calculations. Compounds with the highest binding free energies were selected for further *in vitro* activity assays. The detailed method for FEP-ABFE based virtual screening is given in SI section S1. The derivation of the RED function and extensive evaluations of the accelerated FEP-ABFE method are given in detail in SI section S2 to S7.

### In vitro activity assays of the SARS-CoV-2 M^pro^ inhibitors

The pGEX4T1-M^pro^ plasmid was constructed (AtaGenix, Wuhan) and transfected into the *E. coli* strain BL21 (CodonPlus, Stratagene). A GST-tagged protein was purified by GST-glutathione affinity chromatography and cleaved with thrombin. The purity of the recombinant protein was greater than 95% as assessed by SDS–PAGE. The catalytic activity of M^pro^ was measured by continuous kinetic assays, using an identical fluorogenic substrate MCA-AVLQSGFR-Lys(Dnp)-Lys-NH2 (Apetide Co., Ltd, Shanghai, China). The fluorescence intensity was monitored with a Multifunctional Enzyme Marker (SpectraMax®i3x, Molecular Devices, U.S.A.) using wavelengths of 320 and 405 nm for excitation and emission, respectively. The experiments were performed in a 100 μL reaction system with a buffer consisting of 50 mM Tris-HCl (pH 7.3), 1 mM EDTA. To determine IC_50_ for each compound, the compound was diluted in 100% DMSO to the desired concentrations, solution containing M^pro^ (at the final concentration of 500 nM) was dispensed into black 96-well plates with glass-bottom (Jing’an, Shanghai, China) and was incubated with 1 μL compound at room temperature for 10 min. The reaction was initiated by adding the substrate (at the final concentration of 20 μM). Fluorescence was monitored once every 45 s. Initial reaction velocities were calculated by fitting the linear portion of the curves (within the first 5 min of the progress curves) to a straight line using the program SoftMax Pro and were converted to enzyme activity (substrate cleaved)/second.

## DECLARATION OF INTERESTS

The authors declare no competing financial interest.

## Supporting Information Availability

Additional computational details, computational data, and experimental (including Sections S1 to S7, Figures S1 to S10, and Tables S1 to S6).

## Competing interests

There is no conflict of interests for all authors.

## Acknowledgments

We cordially acknowledge Tencent Cloud and National Supercomputing centers in Shenzhen, Tianjing, and Guangzhou for providing HPC resources for virtual screening and free energy perturbation calculations. We also acknowledge the Beijing Super Cloud Computing Center (BSCC) for providing HPC resources that have contributed to the research results reported within this paper. URL: http://www.blsc.cn/. We cordially acknowledge National Key R&D Program of China (2017YFB0202600), National Natural Science Foundation of China (81903542, 81522041, 21877134), Science Foundation of Guangdong Province (2018A030313215 and 201904020023), Guangdong Provincial Key Laboratory of Construction Foundation (2017B030314030), Fundamental Research Funds for the Central Universities (Sun Yat-Sen University, 18ykpy23), Local Innovative and Research Teams Project of Guangdong Pearl River Talents Program (2017BT01Y093), the National Science and Technology Major Projects for “Major New Drugs Innovation and Development” (2018ZX09711003-003-005), the Strategic Priority Research Program of the Chinese Academy of Sciences (XDC01040100), the National Science Foundation (NSF, grant CHE-1111761), the Taishan Scholars Program (tsqn201909170), the Innovative Leader of Qingdao Program (19-3-2-26-zhc), the special scientific research fund for COVID-19 from the Pilot National Laboratory for Marine Science and Technology (QNLM202001), Sun Yat-Sen University and Zhejiang University special scientific research fund for COVID-19 prevention and control, and philanthropy donation from individuals. The funders had no roles in the design and execution of the study.

## Notes

### Competing Interest Statement

The authors have declared no competing interest.

