## Supporting information for "Identify potent SARS-CoV-2 main protease inhibitors via accelerated free energy perturbation-based virtual screening of existing drugs"

#### Section S1. Detailed method for FEP-ABFE based virtual screening

### S1.1 Molecular docking

The crystal structure (PDB ID: 6LU7)(1) of SARS-CoV-2 $M^{pro}$ which causes COVID-19 was used for molecular docking. Based on the crystal structure, more than 2500 small molecules in the existing drug library (including all FDA-approved drugs) were screened first by using the Glide molecular docking program,(2) followed by the accelerated FEP-ABFE calculations (see below). Considering that the $M^{pro}$ is a protease, Cys145-His41/Ser144-His163 can act as the nucleophilic agent-acid pair to facilitate the catalytic hydrolysis reaction of the substrate proteins, and Gly143 and Gln166 can form hydrogen bonds with the "CO-NH-Cα-CO-NH-Cα" structure of the backbone in the substrate protein. Thus, these 6 residues were considered critical to successful $M^{pro}$ inhibitor screening. After molecular docking, the binding modes of all the ligands were carefully examined, and top-100 molecules with specific interactions with these key residues and relatively high docking scores were selected for further accelerated FEP-ABFE simulations to predict their binding free energies with $M^{pro}$.

### S1.2 Free energy perturbation (FEP)

**S1.2.1 Preliminary molecular dynamics (MD) simulations.** All 100 ligands selected by molecular docking were further evaluated by FEP calculations carried out using the GROMACS-2019 program.(3, 4) Before the FEP calculations, 4 ns preliminary MD simulations were performed for each receptor-ligand complex to improve the fit of the ligand into the binding pocket. All the ligands are parameterized by the general AMBER force field (GAFF).(5) Restrained electrostatic potential (RESP) charge calculations on the ligands were performed using the Gaussian 03 program(6) at the HF/6-31G* level. The parameters of the protein were provided by the AMBER FF14SB force field.(7) The TIP3P force field model(8) was used for water molecules. The systems were neutralized by adding counter ions ($Na^+$ or $Cl^-$) whenever necessary. The systems were first minimized by using the steepest descent method for 5000 cycles and then heated from 0 to 298 K in an NVT ensemble for 100 ps. The systems were then equilibrated in an NPT ensemble with a weak restraint of 1000 kJ/mol/nm$^2$ for 500 ps followed by a 4-ns unconstrained production MD simulation run. The final snapshot of the MD simulated binding structure was used for the subsequent FEP simulation, and the trajectory of the last 2 ns was analyzed to obtain the parameters for adding restraints between the receptors and ligands.

**S1.2.2 Protocol for automatically adding restraints.** Based on the preliminary MD simulation results, FEP-ABFE calculations were carried out according to the thermodynamic cycle presented in Figure 1. As shown in the thermodynamic cycle, a restraint is added to the receptor (or Rec, which is $M^{pro}$) and ligand (or Lig) for each FEP calculation. The strategy of adding restraints, which was proposed first by Boresch *et al.*(9) and used in this study, consists of one distance, two angles, and three dihedral harmonic potentials with a force constant of 10 kcal/mol/$Å^2$ [$rad^2$]. The contribution of the restraints to the Lig system ($\Delta A_{restr}^{L}$) was calculated analytically, and the contribution of the restraint to the Rec-Lig system ($\Delta A_{restr}^{RL}$) was calculated by the FEP simulation. According to the strategy, three atoms of the ligand and three atoms of the receptor are selected and added to the restraints. To add the restraints at the equilibrium position, an in-house program was coded to automatically detect the required parameters and select the three ligand atoms and the three receptor atoms. For the ligand, the heavy atom closest to the geometric center is selected as the first one; the heavy atom that is most distant from the first atom is selected as the second one; the heavy atom which forms an angle with the first two atoms larger than 90 degrees and is most distant from the first atom is selected as the third one. For the receptor, based on the last 2-ns trajectory of the 4-ns preliminary MD simulation, the distances, angles, and dihedrals between the three ligand atoms and the $C_\alpha$, $C_c$ (carbon of the carboxyl group) and N (N atom of the amino group) atoms of all the residues within 5 Å of the ligand were calculated along the MD trajectory. The $C_\alpha$, $C_c$, and N atoms from the same residue with the most stable (associated with the lowest standard deviation values) distances, angles, and dihedral values were selected as the three receptor atoms. The mean values for distance, angles, and dihedrals were used for adding the restraints between the three ligand atoms and the three receptor atoms.

### S1.3 ABFE calculation

To calculate the ABFE for a ligand with its receptor, the ligand electrostatic and van der Waals interactions are decoupled. In recently published works,(10, 11) 12 λ was used for adding the restraints, 10 λ was used for the decoupling electrostatic interactions, and 20 λ was used for the decoupling vdW interactions, which would be more computationally demanding and, thus, is not suitable for efficient virtual screening.

To make it possible using the FEP-ABFE calculations to rapidly identify $M^{pro}$ inhibitors against COVID-19 from existing drugs, the alchemical pathway was optimized to accelerate the FEP. According to the FEP theory, to calculate the free energy difference $\Delta A$, the probability distribution of the potential energy differences between the adjacent λ, denoted as $P(\Delta U)$, is sampled. In reported studies,(12, 13) $P(\Delta U)$ values are considered to have a Gaussian-like distribution, similar to the steps of the electrostatic interaction and vdW interaction decoupling. However, for the added restraint steps, the $P(\Delta U)$ values do not follow a Gaussian distribution. In this study, we first derived and introduced a restraint energy distribution (RED) function which can be used to reasonably describe the $P(\Delta U)$ of the added restraint steps. By using the automatic restraint-adding program described above and fitting $P(\Delta U)$ values by the RED function, the restraint energy can be accurately estimated by merely using single-step perturbation (λ from 0.0 directly to 1.0), which greatly decreases the computational need necessary for the restraint addition. For the decoupling of electrostatic and vdW interactions, the alchemical pathways that can significantly decrease the number of λ parameters needed to maintain accuracy were also studied. After all these procedures were performed, the FEP-ABFE calculations were greatly accelerated, making the practical application of the FEP-ABFE predictions to virtual screening possible.

After the alchemical pathway was determined, for each window, 5000 cycles of the steepest descent energy minimization were carried out first, and then 100-ps simulations in the NVT ensemble along with Langevin dynamics(14, 15) for temperature coupling were performed to heat the system to

298 K with weak position restraints of 1000 kJ/mol/nm$^2$ applied to the receptor and the ligand heavy atoms. The simulation system was subsequently equilibrated in an NPT ensemble for 500 ps with the position restraints continuously applied, followed by 4-ns unconstrained production simulation. Pressure was coupled using the Parrinello–Rahman pressure coupling scheme.(16) The LINCS constraint algorithm(17) was used only on H-bonds. In all the simulations, the particle mesh Ewald (PME) algorithm(18) was used for the calculation of long-range electrostatic interactions. The $\Delta U$ values were sampled during the unconstrained simulation, and the free energy differences between each window were calculated by using the Bennet acceptance ratio (BAR) method.(19, 20)

Several studies(21-23) have reported that the FEP-ABFE method is relatively accurate for electrically neutral ligands, but when the net charge of the ligand is not 0, a systematic error is encountered. Considering that many of the existing drugs are not electrically neutral, all the molecules evaluated via the FEP-ABFE calculations were grouped by their net charges, and in each group, the molecules with the highest binding free energies were selected for further *in vitro* activity assays.

**Section S2. Overview of the RED function-accelerated the FEP-ABFE approach**

The thermodynamic cycle used for the FEP-ABFE calculation is shown in Figure 1 in the main text. The probability distribution of the energy difference between different windows $P(\Delta U)$ used for calculating $\Delta A_{restr}^{RL}$ can be described by the following restraint energy distribution (RED) function (Eq. S1), with the derivation and detailed discussion of the RED function given in the following section (section S3).

$$P(\Delta U) = \frac{1}{\exp(c\Delta U)^{n_1}} a \cdot b^{\frac{3}{2}} \exp(-b\Delta U) \cdot (\Delta U)^2 + \frac{1}{1 + \left(\frac{d}{\Delta U}\right)^{n_2}} \left( h \frac{1}{\sqrt{\pi}\mu_i} \exp\left( \frac{-(\Delta U - \mu)^2}{2\sigma} \right) \right) \quad \text{(S1)}$$

The RED function can accurately describe the sampled restraint energy distribution $P(\Delta U)$ and greatly increase the convergence for calculating $\Delta A_{restr}^{RL}$. As shown in Figure S1a, the sampled $P(\Delta U)$ (yellow dots) used for adding restraints can be fitted quite well by the RED function (red line). By using 3 targets and 28 ligands as a test set and the automatic restraint-adding program and then fitting the $P(\Delta U)$ with the RED function, the $\Delta A_{restr}^{RL}$ calculated by using one-step perturbation (2 λ with values of 0.0 and 1.0) has an excellent correlation with that calculated by using the previously reported method based on 12-λ perturbation with $R^2 > 0.97$ (see Figure S1b). The energy difference between the two alchemical pathways for all the tested systems was less than 0.5 kcal/mol. In addition to using one-step perturbation for the calculation of $\Delta \mathrm{A}_{\mathrm{restr}}^{\mathrm{RL}}$, the alchemical pathway for calculating $\Delta \mathrm{A}_{\mathrm{annihilation}}$ was also optimized, which decreased the number of λ values needed. A detailed discussion of the accuracy of and rationale for using the RED function is provided in SI sections S4 and S5; details of the results showing the comparison between the one-step perturbation and 12-λ perturbation methods are given in SI section S4; and details of the strategy used for further optimizing the alchemical pathway and calculating $\Delta \mathrm{A}_{\mathrm{annihilation}}$ are given in SI section S6. Twenty-eight receptor-ligand systems were used to test the FEP-ABFE method, and the method showed remarkable accuracy as discussed in SI section S7. On the basis of all these efforts, the FEP-ABFE can be calculated accurately by using just 16 λ values and, thus, the FEP-ABFE calculation can be accelerated significantly without losing the accuracy of ABFE prediction. The computational acceleration has made the practical FEP-ABFE prediction-based virtual screening for drug repurposing feasible for the first time.

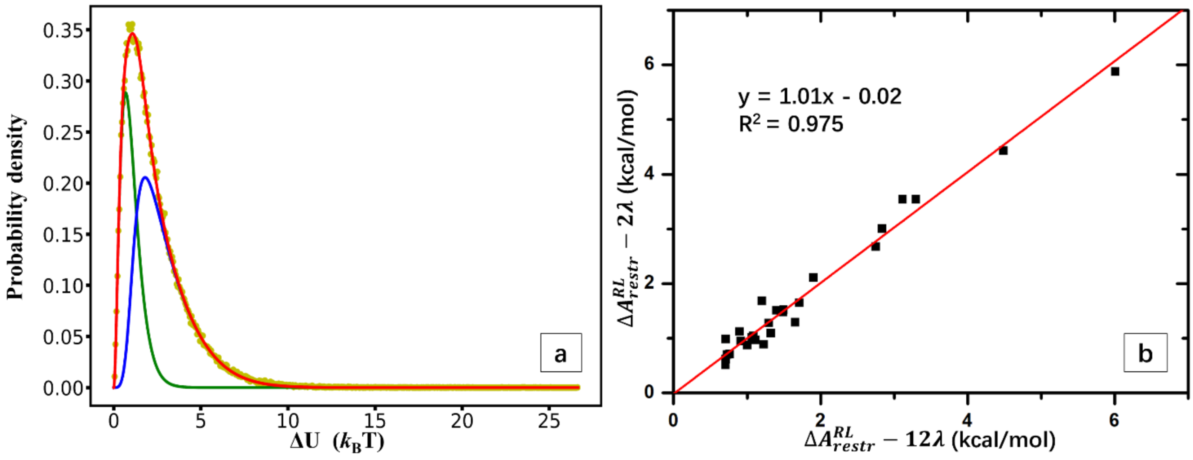


**Figure S1.** (a) P(Δ*U*) can be fitted well with the RED function. The yellow dots are the sampled P(Δ*U*), the red line is the fitted RED function, the green line is the first term in the RED function, and the blue line is the second term in the RED function. (b) The linear regression of the results of $\Delta A_{\text{restr}}^{\text{RL}}$ calculated using the 12-λ perturbation and using the one-step perturbation methods (denoted as 2 λ), which showed an excellent correlation, with a slope of ~1.0, intercept at ~0.0, and $R^2 > 0.97$.

### Section S3. Derivation of the restrain energy distribution (RED) function

Considering the situation where the restraint between the receptor and ligand is added at the equilibrium position, the interaction between receptor and the ligand consists of two parts including the force field and the restraint. Interaction comes from the force field can be simplified as a harmonic potential since the ligand is near the equilibrium, and the restraint force is also a harmonic biasing force. Thus, the total interaction potential between receptor and ligand can be simplified as

$$E_{rec-lig} = k_{total}\,(r - r_0)^2 \qquad \text{(S2)}$$

where $k_{total}$ is the apparent force constant, and $(r - r_0)$ is the generalized distance between the current position and the equilibrium. When adding restraints, the $\Delta U$ between adjacent $\lambda$ will be sampled in order to calculate the free energy of the added restraints, and the $\Delta U$ can be represented by the following equation,

$$\Delta U_{i+1,i} = \Delta\lambda_{i+1,i} k_{res}(r - r_0)^2 \qquad \text{(S3)}$$

where i and i+1 means the adjacent $i$th and $(i + 1)$th window, and $k_{res}$ is the force constant of the added restraint.

Under the above circumstances, the probability distribution $P(\Delta U_{i+1,i})$ can be represented as

$$P(\Delta U_{i+1,i}) = \frac{exp(-\beta E_{rec-lig})\Omega(E_{rec-lig})\Delta U_{i+1,i}}{Z} \qquad \text{(S4)}$$

where Z is the partition function, $\beta$ = $(k_B T)^{-1}$, and $\Omega(E_{rec-lig})$ is the density of state. Since the generalized distance $(r - r_0)$ between the current position $(r)$ and the equilibrium $(r_0)$ can be considered as a three-dimensional vector, the density of state $\Omega(E_{rec-lig})$ can be represented as

$$\Omega(E_{rec-lig}) = 4\pi(r - r_0)^2 \qquad \text{(S5).}$$

Here, in Eq. (S5) we omitted $\frac{V^N}{h^{3N}N!}$ which is a constant factor and will not affect the form of the derived RED function.

By substituting Eqs. (S2), (S3), and (S5) to Eq. (S4), we can get

$$P(\Delta U_{i+1,i}) = \frac{exp(-\beta k_{total}(r - r_0)^2)4\pi(r - r_0)^2\Delta\lambda_{i+1,i}k_{res}(r - r_0)^2}{Z} \qquad \text{(S6),}$$

and by substituting $(r - r_0)^2$ term in Eq. (S6) with $\Delta U_{i+1,i}$ , representing$\Delta U_{i+1,i}$ by $\Delta U$, combining the terms which are constant factors, and normalizing the distribution function, the distribution for $P(\Delta U)$ near the equilibrium has the form of Eq. (S7)

$$P(\Delta U) = a \cdot b^{\frac{3}{2}} \exp(-b\Delta U) \cdot (\Delta U)^2 \qquad (S7),$$

where $a$ and $b$ are all constants, and $b^{\frac{3}{2}}$ is a factor for normalization purpose.

During the simulation, when some of the sampled points are far from the equilibrium, we need to add a Gaussian term to Eq. (S7) to represent such situations, and the $P(\Delta U)$ can be represented as $P(\Delta U) = \frac{1}{\exp(c\Delta U)^{n_1}} a \cdot b^{\frac{3}{2}} \exp(-b\Delta U) \cdot (\Delta U)^2 + \frac{1}{1+\left(\frac{d}{\Delta U}\right)^{n_2}} \left(h \frac{1}{\sqrt{\pi}\mu_i} \exp\left(\frac{-(\Delta U-\mu)^2}{2\sigma}\right)\right) \quad (S8),$

which is the restrain energy distribution (RED) function used in this study, where $a, b, c, d, h, \mu$, and σ are the parameters to be fitted. In the RED function, $\frac{1}{\exp(c\Delta U)^{n_1}}$ and $\frac{1}{1+\left(\frac{d}{\Delta U}\right)^{n_2}}$ are two factors used to combine the two terms and keep the function to be described only by the first term (denoted as harmonic energy term hereinafter) when $\Delta U$ is small and only by the second term (denoted as Gaussian term hereinafter) when $\Delta U$ is large. The two constants $n_1$ and $n_2$ can be set to two relatively big integers, and we chose 10 and 4 for $n_1$ and $n_2$, respectively. One can also choose other integers for $n_1$ and $n_2$ from 2 to 10, that are not expected to affect the fitting results too much. The results of fitting $P(\Delta U)$ by the RED function is discussed in detail in the following section (SI Section S4).

### Section S4. Fitting by the RED function greatly increased the convergence of restraints steps and reduced the calculations needed in the FEP-ABFE prediction

Although the accuracy of the FEP calculations has been reported by several studies,(10, 11) the computational need necessary for a single FEP-ABFE prediction is massive. Thus, in order to rapidly discover inhibitors for clinical use against COVID-19, it is important to increase the convergence and decrease the calculation costs while keeping the accuracy of the FEP simulation. Normally, more than 10 λ values will be calculated during the addition of restraints, such as the works recently reported by Aldeghi *et al.* that 12 non-uniformly distributed λ values are used (0.0, 0.01, 0.025, 0.05, 0.075, 0.1,

0.15, 0.2, 0.3, 0.5, 0.75, 1.0) for the addition of restraints.(10, 11) However, with the use of the automatic restrain adding program and fitting the probability distribution of the sampled energy difference $P(\Delta U)$ function, the $P(\Delta U)$ can be fitted quite well, and the convergence for calculating restraint energy was greatly improved. As a result, the restraint energy can be calculated accurately using just one-step perturbation with λ values change directly from 0.0 to 1.0, and the calculation can be accelerated greatly. Using a HIV-1 protease-ligand complex (crystal structure PDB ID: 2QHY) as an example, the restraint was added by using one-step perturbation and 12-λ perturbation, respectively. The sampled $P(\Delta U)$'s for adding restraints are fitted by the RED function, and the results are shown in Figure S2. In each picture, the yellow dots are the corresponding sampled $P(\Delta U)$, the green line is the harmonic energy term in the RED function, and the blue line is the Gaussian term in the RED function. As expected, the RED function is described only by the harmonic energy term when $\Delta U$ is small and only by the Gaussian term when $\Delta U$ is large, and the sampled $P(\Delta U)$ can be fitted well. The free energies of adding restraints are 0.912 and 0.948 kcal/mol for the 12-λ perturbation and the one-step perturbation, respectively, which means the convergence for the one-step perturbation is pretty good.

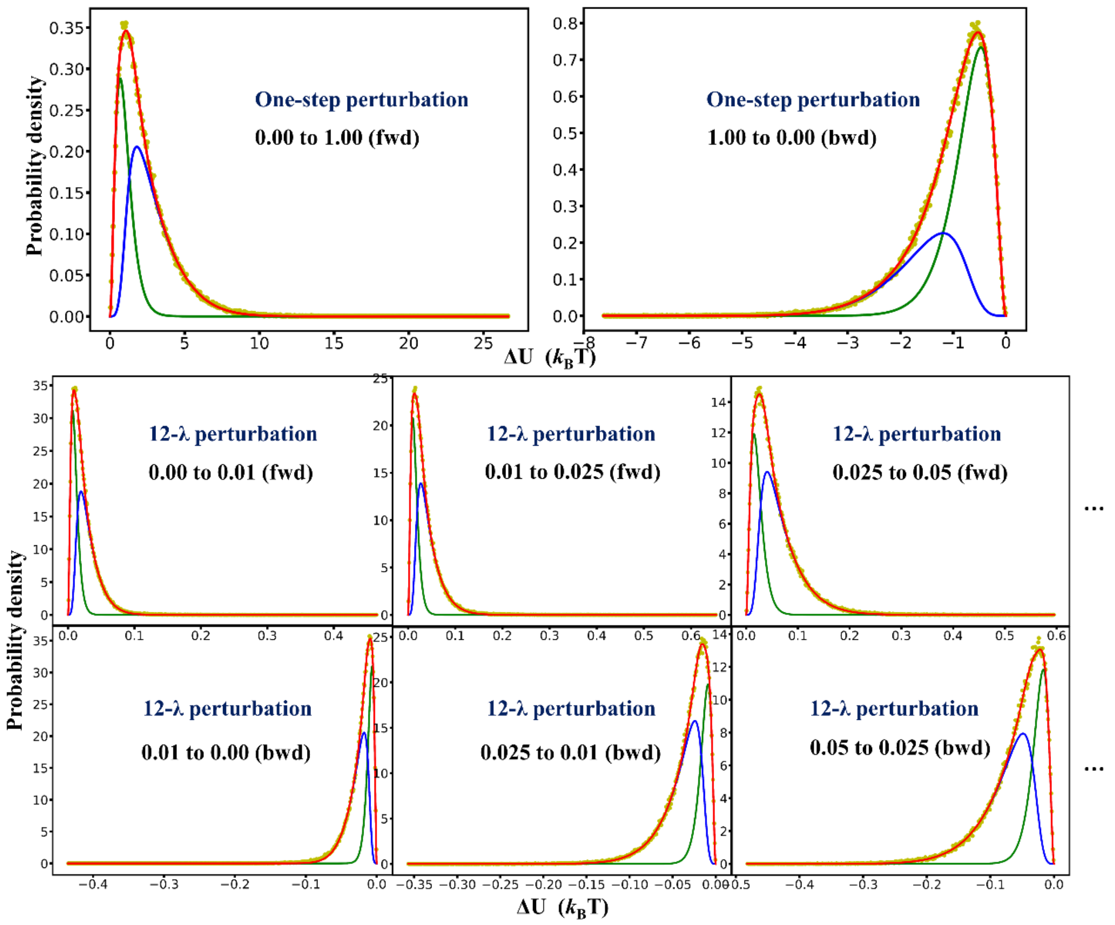


**Figure S2.** The sampled $P(\Delta U)$'s fitted by the RED function. The yellow dots are the corresponding sampled $P(\Delta U)$, the green line is the harmonic energy term in the RED function, and the blue line is the Gaussian term in the RED function.

In order to further justify the applicability of one-step perturbation (2 λ with values of 0.0 and 1.0, denoted as 2 λ below) in adding restraints, three different targets and 28 ligands were tested to compare the $\Delta A_{restr}^{RL}$ results calculated for 12 λ and 2 λ. The bond, angle, and dihedral parameters for the restraints are the same for 12 λ and 2 λ calculations. Table S1 summarized the results for the $\Delta A_{restr}^{RL}$, and the calculation results revealed that the single-step perturbation has the remarkable convergence with the energy difference for all the tested systems less than 0.5 kcal/mol. Thus, by using automatic restrain adding program and fitting the sampled $P(\Delta U)$ by the RED function, the computational need necessary for adding restraints can be greatly decreased without losing the accuracy.

**Table S1.** Comparison of $\Delta A^{RL}_{restr}$ between calculations using 12 λ values and 2 λ values.[a]

| Protein targets | PDB | $\Delta A^{RL}_{12restr}$ [b] | $\Delta A^{RL}_{2restr}$ [c] | $\Delta\Delta A^{RL}_{restr}$ [d] |
|---|---|---|---|---|
| HIV-1 protease | 2QHY | 0.912 | 0.948 | -0.036 |
| | 4U8W | 1.486 | 1.481 | +0.006 |
| | 5UPZ | 1.323 | 1.097 | +0.226 |
| | 1AJV | 0.708 | 0.989 | -0.281 |
| | 1D4H | 4.483 | 4.431 | +0.052 |
| | 1D4I | 0.897 | 1.127 | -0.230 |
| | 1EBY | 1.222 | 0.891 | +0.331 |
| | 1EBZ | 0.712 | 0.631 | +0.080 |
| | 3A2O | 0.706 | 0.513 | +0.192 |
| | 1G2K | 6.009 | 5.878 | +0.131 |
| Human factor Xa | 1F0S | 1.900 | 2.113 | -0.214 |
| | 1FJS | 1.652 | 1.296 | +0.356 |
| | 1MQ6 | 1.494 | 1.520 | -0.026 |
| | 1NFW | 3.296 | 3.545 | -0.301 |
| | 1NFX | 1.706 | 1.651 | +0.055 |
| | 2J34 | 2.747 | 2.681 | +0.066 |
| | 2P16 | 0.725 | 0.703 | +0.021 |
| | 2P95 | 0.710 | 0.622 | +0.048 |
| | 2XC0 | 1.111 | 0.972 | +0.139 |
| | 2VVV | 3.114 | 3.545 | -0.431 |
| BRD4 | 3MXF | 1.197 | 1.687 | -0.490 |
| | 4MR3 | 1.398 | 1.514 | -0.116 |
| | 3U5L | 1.002 | 0.876 | +0.126 |
| | 4MR4 | 0.761 | 0.716 | +0.045 |
| | 3U5J | 1.082 | 1.049 | +0.034 |
| | 3SVG | 1.068 | 1.023 | +0.301 |
| | 4HBV | 1.294 | 1.282 | +0.013 |
| | 4J0R | 2.839 | 3.009 | -0.170 |

[a] The unit of the free energy is kcal/mol.

[b] $\Delta A^{RL}_{12restr}$ is the restraint energy calculated by using 12 λ values and BAR method.

[c] $\Delta A^{RL}_{2restr}$ is the restraint energy calculated from one step perturbation with the use of 2 λ value (0.0, 1.0). The sampled energy distribution from each λ was first fitted by RED function, and then BAR method was used to calculate the energy estimates.

[d] $\Delta\Delta A^{RL}_{restr}$ is the difference of $\Delta A^{RL}_{12restr}$and $\Delta A^{RL}_{2restr}$.

**Section S5. The RED function can also reasonably describe the restraint energy distribution even if the restraint is not added to the equilibrium position**

Although the automatic restrain adding program can make sure that the parameters chosen for the restraint are close to those in the equilibrium, it is still important to have an understanding about the behavior of the probability distribution of $\Delta U$ when the parameters are not at the equilibrium. As an example, we determined the restraint parameters for 2QHY by using the original crystal structure instead by using the pre-equilibrated trajectory, and the resulted $P(\Delta U)'s$ are shown in Figure S3. As seen in Figure S3, the probability distributions for the initial several windows showed distributions with "double-peak" shapes. In theory, the "double-peak" shape distribution $P(\Delta U)$ will happen when relative position between the receptor and ligand, which we use to determine the ligand restraints addition parameters, is far from the equilibrium state, and at the same time, the harmonic potentials used to apply the restraint are not strong enough. According to the FEP theory, the probability distribution near the lower tail of $P(\Delta U)$ will greatly influence the accuracy of $\Delta A$ calculation. The "double-peak" shape of the probability distribution is far from the equilibrium, and the distribution pattern near the lower tail varies from time to time during the simulation, which will result in a very unstable free energy estimation. Thus, to deal with the instability in energy calculation, in the previously reported studies,(10, 11) the restraints are added with more than 10 λ values, and the first several λ are selected rather close to each other (0.0, 0.01, 0.025, 0.05, 0.075, 0.1). Although the purpose of the RED function is to describe the $P(\Delta U)$ when the restraints are added near the equilibrium state, the RED function can also describe the $P(\Delta U)$ well even when restraints are not added near the equilibrium. Using the forward perturbation from 0.00 to 0.01 as an example, which is shown in Figure S3, when the added restraint is relatively weak and not at the equilibrium position, the peak on the right side (which is described by the Gaussian term of the RED function) is caused by the ligand occupying the equilibrium position, and the peak on the left is caused by the added restraint described by the harmonic energy term of the RED function. The RED function can describe the $P(\Delta U)$ of adding restraints quite well no matter whether the restraints are added near the equilibrium, which further proved the correctness and rationality of the function. The RED function can be used to further optimize, and give a deeper understanding of, the FEP-ABFE calculation procedure in the future studies.

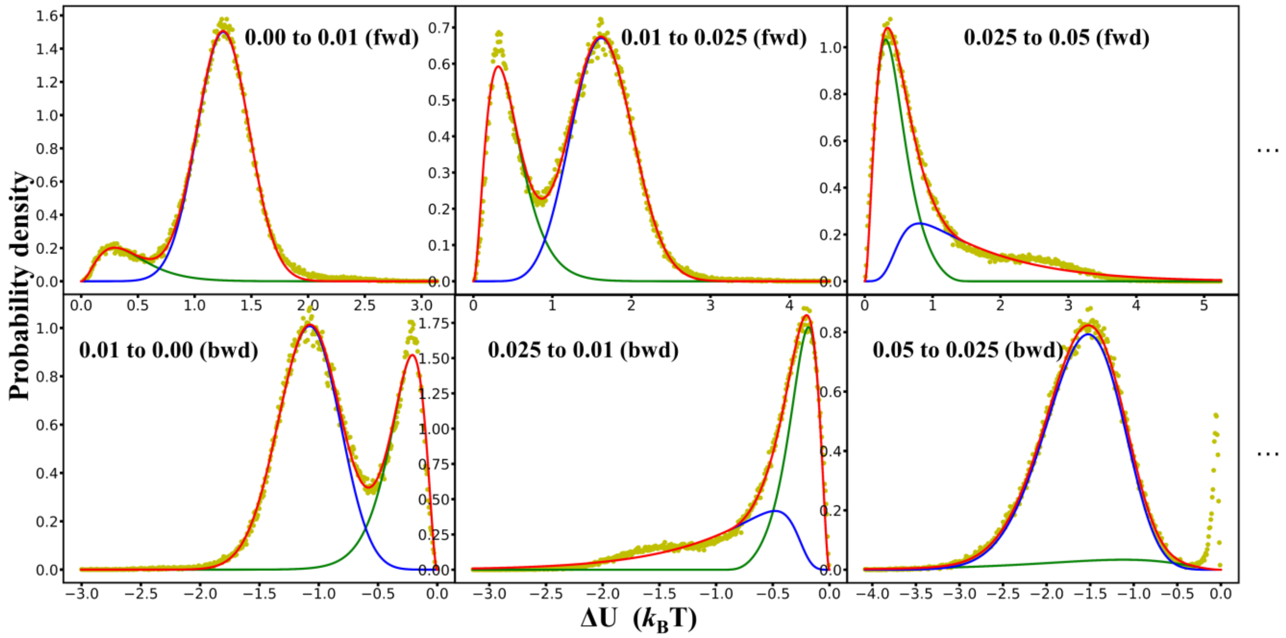


**Figure S3.** The sampled $P(\Delta U)$'s fitted by the RED function when the restraints are not added near the equilibrium position. The yellow dots are the corresponding sampled $P(\Delta U)$, the green line is the harmonic energy term in the RED function, and the blue line is the Gaussian term in the RED function.

### Section S6. FEP calculations can be accelerated by more than ~3-fold without losing the accuracy with appropriate selection of λ values

In order to further accelerate the calculation, we studied the effect of decreasing the number of λ values for both charge and vdW interactions decoupling by using 4 different receptor-ligand systems. Corresponding crystal structures (PDB codes: 3SVG, 3U5L, 3U5J, 4HBV) were used in order to make sure that the binding modes between the receptor and ligands are reasonable. For all the 4 crystal structures, $\Delta A^{RL}$ was first calculated by using 42 λ (12 λ for restrain, 10 λ for charge, 20 λ for vdW) as reported by the recently published works,(10, 11) and then the number of λ values were decreased for the decoupling of both charge and vdW interactions. As seen in Table S2, by using 16 λ values in total (2 λ for restrain with fitting by RED function, 5 λ for charge, and 9 λ for vdW) the energy calculation results are quite close to those calculated by using 42 λ values for all the 4 systems tested (all the energy differences are within 1 kcal/mol). Compared with the 42 λ, the FEP-ABFE calculations using just 16 λ values need only 38 % computational resources, and the calculations are accelerated by ~3-fold.

**Table S2.** The details of free energy changes for the complex in the annihilation for the traditional 42-window scheme and the 16-window scheme.[a]

| λ [b] | 3SVG | | 3U5L | | 3U5J | | 4HBV | |
|---|---|---|---|---|---|---|---|---|
| | $\Delta A^{RL}_{42\,\lambda}$ [c] | $\Delta A^{RL}_{16\,\lambda}$ [d] | $\Delta A^{RL}_{42\,\lambda}$ | $\Delta A^{RL}_{16\,\lambda}$ | $\Delta A^{RL}_{42\,\lambda}$ | $\Delta A^{RL}_{16\,\lambda}$ | $\Delta A^{RL}_{42\,\lambda}$ | $\Delta A^{RL}_{16\,\lambda}$ |
| (0.0, 0.0, 0.0) | 0.000 | 0.000 | 0.000 | 0.000 | 0.000 | 0.000 | 0.000 | 0.000 |
| (0.01, 0.0, 0.0) | 0.239 | - | 0.016 | - | 0.029 | - | 0.030 | - |
| (0.025,0.0,0.0) | 0.394 | - | 0.055 | - | 0.067 | - | 0.085 | - |
| (0.05, 0.0, 0.0) | 0.539 | - | 0.117 | - | 0.121 | - | 0.165 | - |
| (0.075,0.0,0.0) | 0.697 | - | 0.154 | - | 0.165 | - | 0.219 | - |
| (0.10, 0.0, 0.0) | 0.914 | - | 0.202 | - | 0.204 | - | 0.271 | - |
| (0.15, 0.0, 0.0) | 1.365 | - | 0.290 | - | 0.277 | - | 0.366 | - |
| (0.20, 0.0, 0.0) | 1.681 | - | 0.346 | - | 0.369 | - | 0.449 | - |
| (0.30, 0.0, 0.0) | 2.086 | - | 0.476 | - | 0.528 | - | 0.592 | - |
| (0.50, 0.0, 0.0) | 2.736 | - | 0.699 | - | 0.723 | - | 0.833 | - |
| (0.75, 0.0, 0.0) | 3.416 | - | 0.863 | - | 0.924 | - | 1.082 | - |
| (1.0, 0.0, 0.0) | 3.992 | 3.242 | 1.002 | 0.912 | 1.082 | 1.115 | 1.294 | 1.311 |
| (1.0, 0.1, 0.0) | 6.963 | - | 3.123 | - | 3.184 | - | 3.715 | - |
| (1.0, 0.2, 0.0) | 9.391 | 9.129 | 5.077 | 4.987 | 5.122 | 5.152 | 5.815 | 5.380 |
| (1.0, 0.3, 0.0) | 11.611 | - | 6.874 | - | 6.881 | - | 7.613 | - |
| (1.0, 0.4, 0.0) | 13.466 | 12.991 | 8.456 | 8.368 | 8.429 | 8.447 | 9.259 | 8.961 |
| (1.0, 0.5, 0.0) | 14.830 | - | 9.817 | - | 9.821 | - | 10.667 | - |
| (1.0, 0.6, 0.0) | 15.896 | 15.339 | 10.999 | 11.050 | 10.978 | 10.975 | 11.613 | 11.248 |
| (1.0, 0.7, 0.0) | 16.669 | - | 12.003 | - | 11.931 | - | 12.253 | - |
| (1.0, 0.8, 0.0) | 17.236 | 16.627 | 12.758 | 12.786 | 12.568 | 12.825 | 12.692 | 12.601 |
| (1.0, 0.9, 0.0) | 17.631 | - | 13.285 | - | 13.012 | - | 12.902 | - |
| (1.0, 1.0, 0.0) | 17.857 | 17.198 | 13.662 | 13.650 | 13.526 | 13.530 | 12.985 | 13.166 |
| (1.0, 1.0, 0.05) | 19.052 | - | 15.025 | - | 14.877 | - | 13.992 | - |
| (1.0, 1.0, 0.10) | 20.241 | 19.606 | 16.387 | 16.401 | 16.205 | 16.075 | 14.995 | 15.157 |
| (1.0, 1.0, 0.15) | 21.395 | - | 17.738 | - | 17.503 | - | 15.979 | - |
| (1.0, 1.0, 0.20) | 22.505 | 21.945 | 19.070 | 19.096 | 18.737 | 18.590 | 16.977 | 17.198 |
| (1.0, 1.0, 0.25) | 23.599 | - | 20.381 | - | 19.928 | - | 17.978 | - |
| (1.0, 1.0, 0.30) | 24.631 | 24.187 | 21.657 | 21.669 | 21.107 | 20.934 | 18.969 | 19.225 |
| (1.0, 1.0, 0.35) | 25.617 | - | 22.914 | - | 22.247 | - | 19.939 | - |
| (1.0, 1.0, 0.40) | 26.622 | - | 24.170 | - | 23.323 | - | 20.889 | - |
| (1.0, 1.0, 0.45) | 27.580 | - | 25.419 | - | 24.365 | - | 21.847 | - |
| (1.0, 1.0, 0.50) | 28.516 | 28.207 | 26.652 | 26.735 | 25.433 | 25.271 | 22.825 | 23.204 |
| (1.0, 1.0, 0.55) | 29.379 | - | 27.876 | - | 26.487 | - | 23.790 | - |
| (1.0, 1.0, 0.60) | 30.118 | - | 28.984 | - | 27.458 | - | 24.713 | - |
| (1.0, 1.0, 0.65) | 30.660 | - | 30.000 | - | 28.361 | - | 25.501 | - |
| (1.0, 1.0, 0.70) | 31.043 | 30.754 | 30.883 | 31.052 | 28.974 | 28.881 | 25.961 | 26.450 |
| (1.0, 1.0, 0.75) | 31.292 | - | 31.599 | - | 29.356 | - | 26.174 | - |

| | | | | | | | | |
|---|---|---|---|---|---|---|---|---|
| (1.0, 1.0, 0.80) | 31.108 | 30.866 | 32.084 | 32.223 | 29.559 | 29.733 | 26.115 | 26.389 |
| (1.0, 1.0, 0.85) | 30.272 | - | 31.945 | - | 29.208 | - | 25.673 | - |
| (1.0, 1.0, 0.90) | 29.402 | 29.477 | 31.368 | 31.196 | 28.490 | 29.334 | 25.189 | 25.507 |
| (1.0, 1.0, 0.95) | 28.986 | 29.101 | 31.069 | 30.670 | 28.098 | 28.929 | 24.969 | 25.329 |
| (1.0, 1.0, 1.0) | 29.156 | 29.259 | 31.237 | 30.728 | 28.206 | 29.019 | 25.128 | 25.468 |

[a] All the free energy differences are calculated by the BAR method. The unit of free energy differences is kcal/mol.

[b] In the lambda arrays, the first value refers to the restraint-lambdas and the second and third value represents the value of coul-lambdas and vdw-lambdas.

[c] $\Delta A^{RL}_{42\,\lambda}$ refers to the free energy difference between the initial state of perturbation and the specific alchemical state defined by the relative lambda arrays, whose alchemical pathway is defined by 42 windows.

[d] $\Delta A^{RL}_{16\,\lambda}$ refers to the free energy difference between the initial state of perturbation and the specific alchemical state defined by the relative lambda arrays, whose alchemical pathway is defined by 16 windows.

### Section S7. Remarkable accuracy of the FEP-ABFE calculations based on the test results from 28 receptor-ligand systems

The accuracy of the accelerated 16-λ-FEP-ABFE calculation was first tested against 3 targets with 28 receptor-ligand systems. One of the targets was BRD4, which was also the focus of a study of the FEP-ABFE method using 42 λ values previously reported by Aldeghi *et al.*,(10, 11) and since the FEP-ABFE calculations show systematic bias when the net charge of a ligand is not 0, all 8 neutral ligands from the work of Aldeghi *et al*. were used in this study.(10) In addition to the target BRD4, the other two targets were HIV-1 protease (with 10 electrically neutral ligands) and human factor Xa (with 10 electrically neutral ligands). All the binding modes for the 28 calculated test systems were obtained from the protein data bank (www.rcsb.org), which ensured the correctness of the initial structures. As shown in detail in Figures S5-S7 and Table S3, the ligands were quite diverse in terms of the following features: the molecular weights ranged from 241 to 662 Da; number of atoms ranged from 22 to 89; number of rotatable bonds ranged from 0 to 21; number of hydrogen bond acceptors ranged from 1 to 7; number of hydrogen bond donors ranged from 0 to 6; and the calculated log*P* ranged from 1.17 to 4.85. The FEP-ABFE calculation results for all the 28 systems are summarized in Table S4, and the linear regression statistics for the calculated and experimental results are given in Figure S4. For the HIV-1 protease and human factor Xa, most of the predicted binding free energy ($\Delta G_{pred}$) values are consistent with the corresponding experimental binding free energy ($\Delta G_{exp}$) values, with an average prediction error of less than 2.0 kcal/mol. For BRD4, although the calculations showed some systematic error, with all the calculation results shifting in the negative direction, the calculation results still showed a good linear correlation with the experimental results. For comparison, the commonly used binding free energy calculation methods MM-PBSA (molecular mechanics Poisson-Boltzmann surface area) and MM-GBSA (molecular mechanics generalized-Born surface area) were also used to calculate the binding free energies for all the 28 receptor-ligand systems, with the detailed results given in Table S5 and Figure S4.

As seen in Figure S4, most of the results from the MM-PBSA and MM-GBSA calculations showed very poor correlation (or no significant correction) with the experimental data, with $R^2 = 0.000$ to 0.136 for the MM-PBSA results and $R^2 = 0.150$ to 0.366 for the MM-GBSA results for the three targets. In comparison, $R^2 = 0.642$ to 0.915 for the FEP-ABFE results, as seen in Figure S4. So, the FEP-ABFE results for all the three targets are markedly better than the corresponding MM-PBSA and MM-GBSA results. The test results based on the 28 ligands with diverse chemical scaffolds suggested that the accelerated FEP-ABFE algorithm can achieve a remarkable accuracy, which encouraged us to perform the FEP-ABFE prediction-based practical virtual screening to identify SARS-CoV-2 $M^{pro}$ inhibitors for drug repurposing.

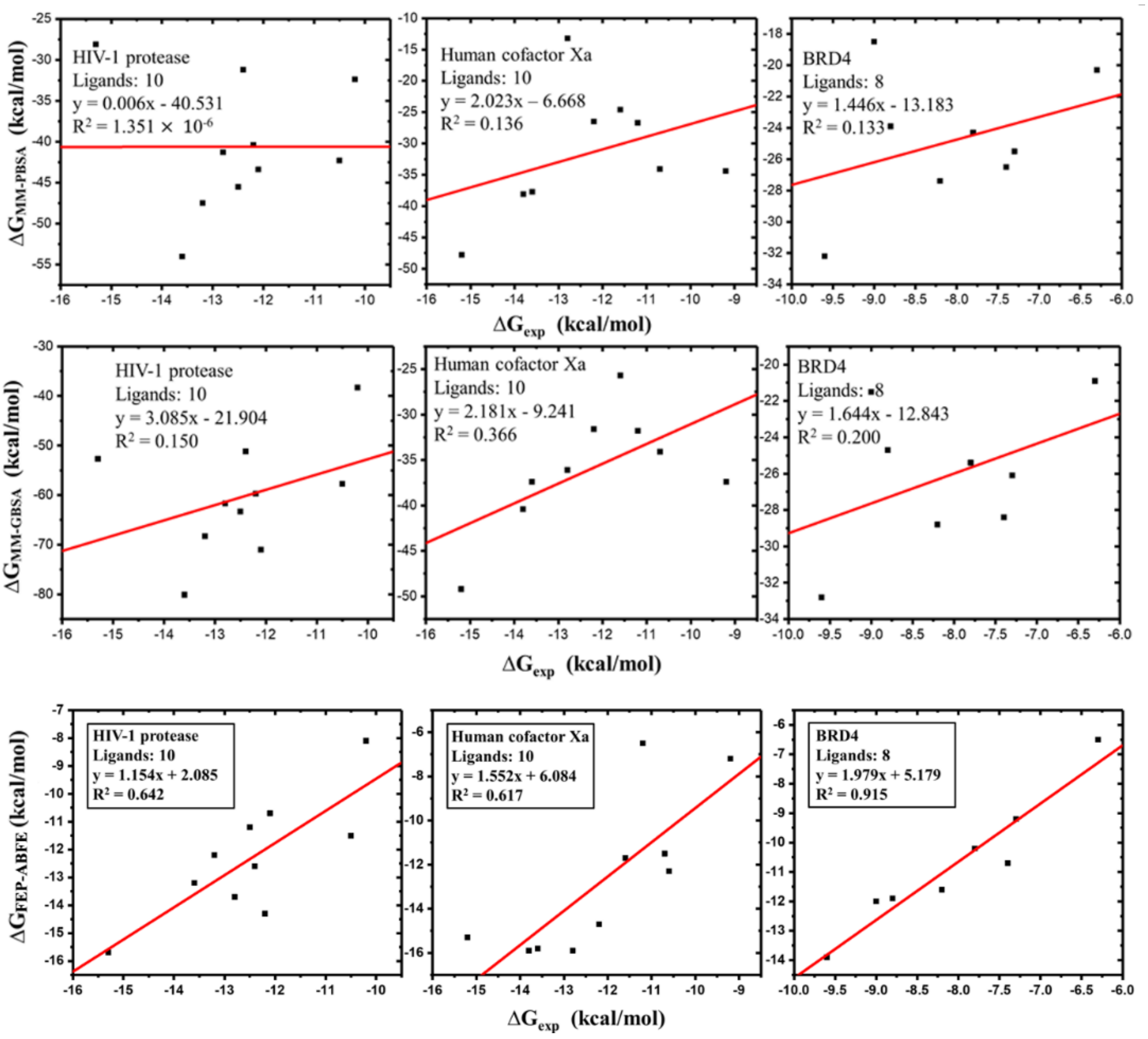


**Figure S4.** The regression models between experimental $\Delta G_{exp}$ and predicted $\Delta G_{MM\text{-}PBSA}$, $\Delta G_{MM\text{-}GBSA}$, and $\Delta G_{FEP\text{-}ABFE}$ values for the three targets and 28 ligands.

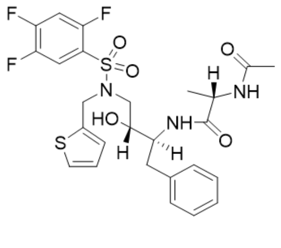


2QHY

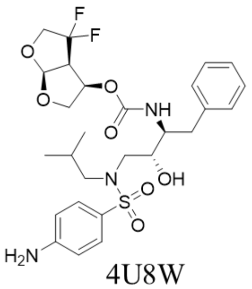


4U8W

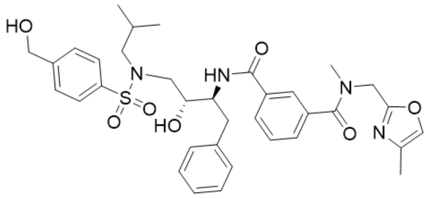


5UPZ

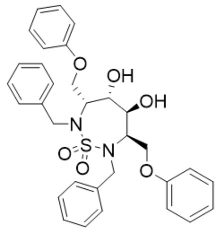


1AJV

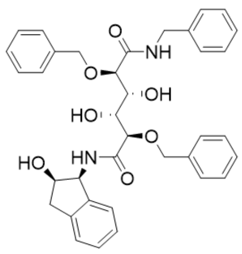


1D4H

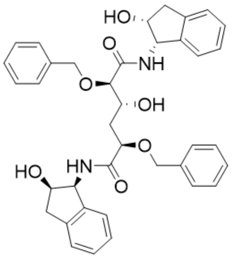


1D4I

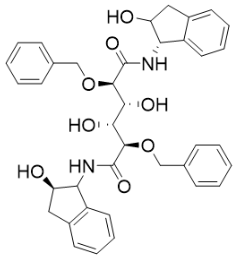


1EBY

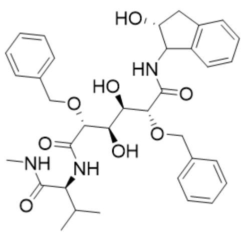


1EBZ

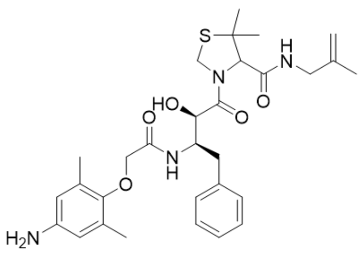


3A2O

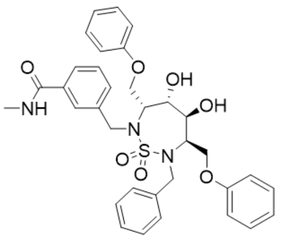


1G2K

**HIV-1 protease**

**Figure S5.** Molecular structures of the tested HIV-1 protease inhibitors

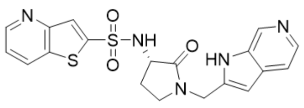

1F0S

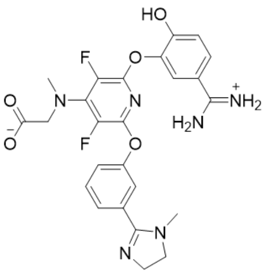

1FJS

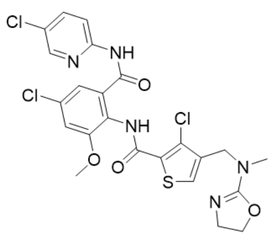

1MQ6

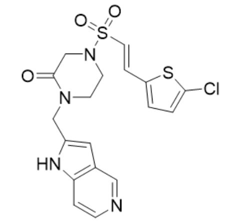

1NFW

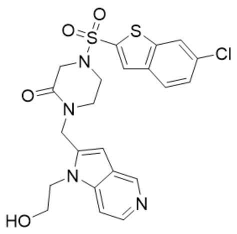

1NFX

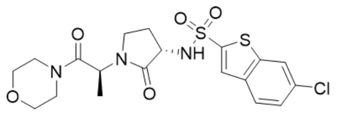

2J34

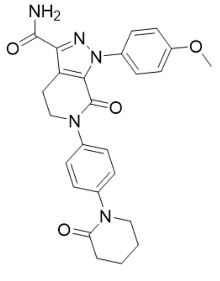

2P16

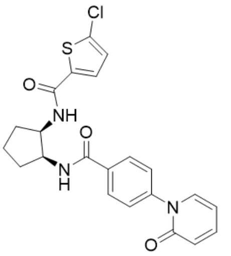

2P95

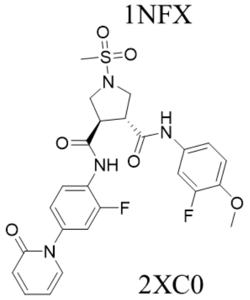

2XC0

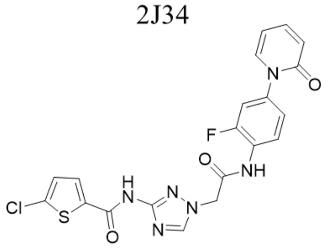

2VVV

**Human cofactor Xa**

**Figure S6.** Molecular structures of the tested human cofactor Xa inhibitors

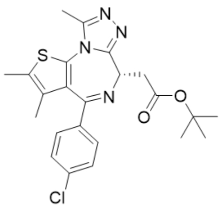

3MXF

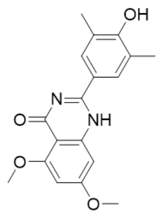

4MR3

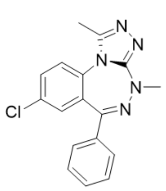

3U5L

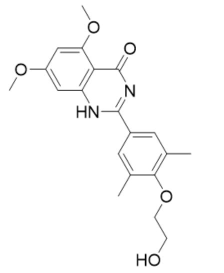

4MR4

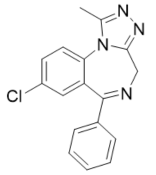

3U5J

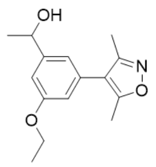

3SVG

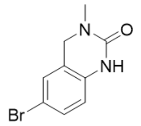

4HBV

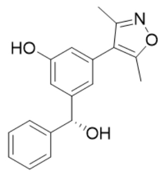

4J0R

**BRD4**

**Figure S7.** Molecular structures of the tested BRD4 inhibitors.

**Table S3.** Physical chemical properties of the 28 tested ligands. PDB refers the PDB code with bound ligands. MW is the molecular weight in Daltons; Netchg is the net charge; NROT is the number of rotatable bonds; HBA is the number of hydrogen bond acceptors; HBD is the number of hydrogen bond donors; cLogP is the calculated octanol/water partition coefficient (computed with XLOGP3). All the properties are gained from the PDBbind-CN Database.

| PDB | MW(Da) | No. Atoms | Netchg | NROT | HBA | HBD | cLogP |
|---|---|---|---|---|---|---|---|
| 2QHY | 583.643 | 67 | 0 | 15 | 5 | 3 | 3.05 |
| 4U8W | 583.645 | 75 | 0 | 14 | 4 | 3 | 3.25 |
| 5UPZ | 662.796 | 89 | 0 | 19 | 7 | 3 | 3.70 |
| 1AJV | 574.687 | 75 | 0 | 12 | 4 | 2 | 4.42 |
| 1D4H | 610.696 | 83 | 0 | 19 | 5 | 5 | 3.43 |
| 1D4I | 636.284 | 87 | 0 | 18 | 5 | 5 | 3.86 |
| 1EBY | 652.733 | 88 | 0 | 19 | 6 | 6 | 3.44 |
| 1EBZ | 633.731 | 89 | 0 | 21 | 6 | 6 | 2.65 |
| 3A2O | 568.727 | 80 | 0 | 15 | 4 | 4 | 3.97 |
| 1G2K | 631.738 | 82 | 0 | 14 | 5 | 3 | 3.69 |
| 1F0S | 427.500 | 46 | 0 | 5 | 5 | 2 | 1.54 |
| 1FJS | 526.500 | 63 | 0 | 10 | 4 | 1 | 4.76 |
| 1MQ6 | 568.860 | 56 | 0 | 10 | 3 | 2 | 4.66 |
| 1NFW | 436.936 | 45 | 0 | 5 | 4 | 1 | 2.06 |
| 1NFX | 505.010 | 54 | 0 | 7 | 5 | 1 | 2.14 |
| 2J34 | 471.978 | 52 | 0 | 6 | 4 | 1 | 2.12 |
| 2P16 | 459.497 | 59 | 0 | 5 | 4 | 1 | 2.24 |
| 2P95 | 441.931 | 50 | 0 | 7 | 3 | 2 | 4.15 |
| 2XC0 | 546.543 | 62 | 0 | 9 | 5 | 2 | 1.17 |
| 2VVV | 472.880 | 46 | 0 | 8 | 5 | 2 | 3.28 |
| 3MXF | 456.988 | 56 | 0 | 5 | 3 | 0 | 4.85 |
| 4MR3 | 326.347 | 42 | 0 | 4 | 2 | 2 | 2.67 |
| 3U5L | 323.780 | 37 | 0 | 1 | 3 | 0 | 4.57 |
| 4MR4 | 370.399 | 49 | 0 | 7 | 2 | 2 | 2.31 |
| 3U5J | 308.765 | 35 | 0 | 1 | 2 | 0 | 3.23 |
| 3SVG | 261.321 | 38 | 0 | 4 | 4 | 1 | 2.21 |
| 4HBV | 241.085 | 22 | 0 | 0 | 1 | 1 | 1.37 |
| 4J0R | 295.332 | 39 | 0 | 5 | 3 | 2 | 3.25 |

**Table S4.** Summary of the FEP free energy calculation results (kcal/mol) for the three important drug targets including HIV-1 protease, human factor Xa, and BRD4.

| Target | PDB | $\Delta G_{exp}$ | $\Delta G_{cal}$ | $\Delta G_{cal} - \Delta G_{exp}$ |
|---|---|---|---|---|
| HIV-1 protease | 2QHY | -10.2 | -8.1 | +2.1 |
| | 4U8W | -15.3 | -15.7 | -0.4 |
| | 5UPZ | -12.5 | -11.2 | +1.3 |
| | 1AJV | -10.5 | -11.5 | -0.9 |
| | 1D4H | -13.6 | -13.2 | +0.4 |
| | 1D4I | -12.1 | -10.7 | +1.3 |
| | 1EBY | -13.2 | -12.2 | +1.0 |
| | 1EBZ | -12.8 | -13.7 | -0.8 |
| | 3A2O | -12.4 | -12.6 | -0.3 |
| | 1G2K | -12.2 | -14.3 | -2.1 |
| Human factor Xa | 1F0S | -10.6 | -12.3 | -1.7 |
| | 1FJS | -13.6 | -15.8 | -2.2 |
| | 1MQ6 | -15.2 | -15.3 | -0.1 |
| | 1NFW | -12.2 | -14.7 | -2.5 |
| | 1NFX | -11.6 | -11.7 | -0.1 |
| | 2J34 | -10.7 | -11.5 | -0.9 |
| | 2P16 | -13.8 | -15.9 | -2.1 |
| | 2P95 | -12.8 | -15.9 | -3.2 |
| | 2XC0 | -9.2 | -7.2 | +1.9 |
| | 2VVV | -11.2 | -6.5 | +4.7 |
| BRD4 | 3MXF | -9.6 | -13.9 | -4.3 |
| | 4MR3 | -9.0 | -12.0 | -3.0 |
| | 3U5L | -8.2 | -11.6 | -3.4 |
| | 4MR4 | -7.8 | -10.2 | -2.4 |
| | 3U5J | -7.4 | -10.7 | -3.3 |
| | 3SVG | -7.3 | -9.2 | -1.9 |
| | 4HBV | -6.3 | -6.5 | -0.2 |
| | 4J0R | -8.8 | -11.9 | -3.1 |

$\Delta G_{exp}$ and $\Delta G_{cal}$ are the experimental binding free energy and FEP-calculated binding free energy, respectively.

**Table S5.** Summary of the MM-PBSA and MM-GBSA calculation results (kcal/mol) for the HIV-1 protease, human factor Xa, and BRD4 inhibitors. Eight nanoseconds MD simulations were performed for each receptor-ligand system, and MM-PBSA and MM-GBSA were calculated based on 100 snapshots extracted from the trajectories of the last 1 ns MD simulations (with an interval of 10 ps).

| Target | PDB | $\Delta G_{exp}$ | $\Delta G_{MM\_PBSA}$ | $\Delta G_{MM\_GBSA}$ |
|---|---|---|---|---|
| HIV-1 protease | 2QHY | -10.2 | -32.4±4.2 | -38.3±3.4 |
| | 4U8W | -15.3 | -28.1±5.7 | -52.7±4.2 |
| | 5UPZ | -12.5 | -45.5±6.0 | -63.3±3.6 |
| | 1AJV | -10.5 | -42.3±4.6 | -57.7±3.8 |
| | 1D4H | -13.6 | -54.0±5.3 | -80.1±4.6 |
| | 1D4I | -12.1 | -43.4±5.9 | -71.0±4.7 |
| | 1EBY | -13.2 | -47.5±5.7 | -68.3±4.4 |
| | 1EBZ | -12.8 | -41.3±5.5 | -61.7±4.3 |
| | 3A2O | -12.4 | -31.2±6.4 | -51.2±4.0 |
| | 1G2K | -12.2 | -40.4±7.5 | -59.7±3.9 |
| Human factor Xa | 1F0S | -10.6 | -22.5±3.5 | -25.4±2.6 |
| | 1FJS | -13.6 | -37.7±8.1 | -37.4±6.8 |
| | 1MQ6 | -15.2 | -47.8±3.2 | -49.2±2.9 |
| | 1NFW | -12.2 | -26.5±3.0 | -31.6±2.4 |
| | 1NFX | -11.6 | -24.6±3.1 | -25.7±2.2 |
| | 2J34 | -10.7 | -34.1±3.1 | -34.1±2.7 |
| | 2P16 | -13.8 | -38.1±3.4 | -40.4±2.7 |
| | 2P95 | -12.8 | -13.2±8.3 | -36.1±2.4 |
| | 2XC0 | -9.2 | -34.4±4.2 | -37.4±4.6 |
| | 2VVV | -11.2 | -26.7±3.1 | -31.8±2.0 |
| BRD4 | 3MXF | -9.6 | -32.2±2.5 | -32.8±1.8 |
| | 4MR3 | -9.0 | -18.5±2.9 | -21.5±1.9 |
| | 3U5L | -8.2 | -27.4±2.6 | -28.8±2.3 |
| | 4MR4 | -7.8 | -24.3±2.6 | -25.4±2.2 |
| | 3U5J | -7.4 | -26.5±2.2 | -28.4±1.9 |
| | 3SVG | -7.3 | -25.5±2.5 | -26.1±2.3 |
| | 4HBV | -6.3 | -20.3±2.4 | -20.9±1.7 |
| | 4J0R | -8.8 | -23.9±2.2 | -24.7±2.0 |

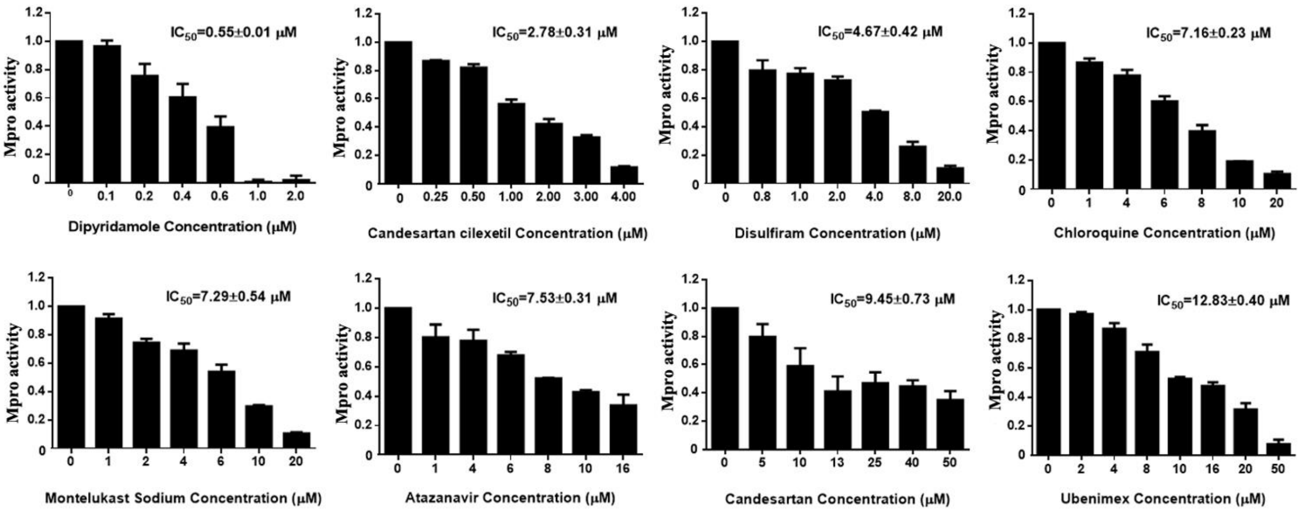


**Figure S8.** The inhibitory curves for the most potent M$^{pro}$ inhibitors.

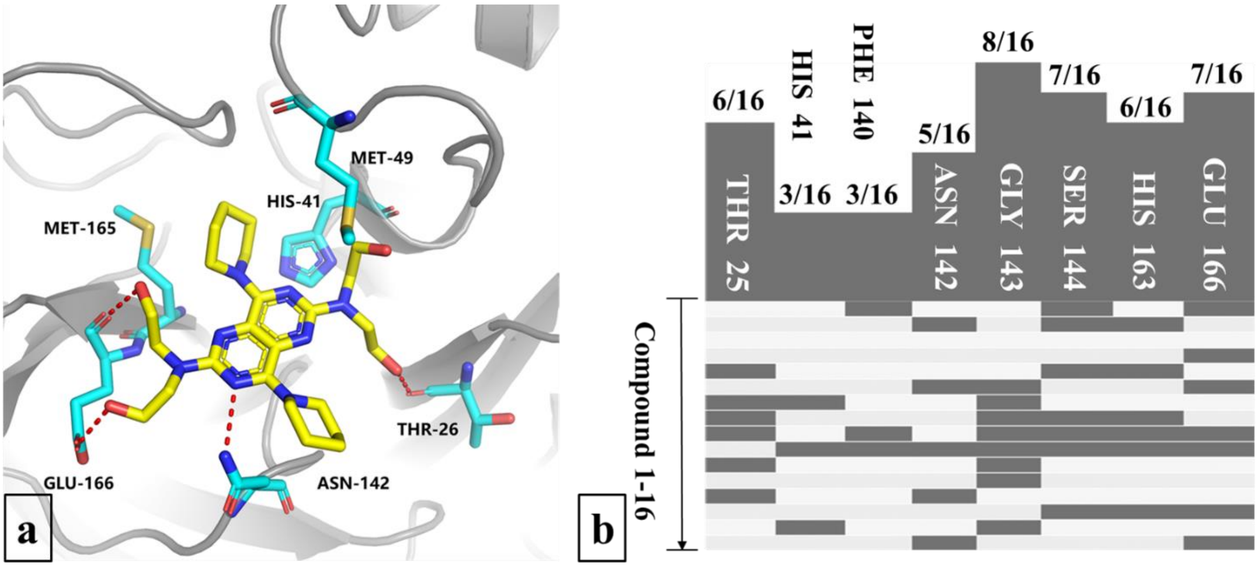


**Figure S9.** (a) Binding mode of M$^{pro}$ with DIP after the MD simulation. DIP is shown in yellow in the stick model, the key amino-acid residues of M$^{pro}$ are shown in cyan in the stick model, and the hydrogen bonds are shown as red dashed lines. (b) The protein-ligand interaction fingerprint (PLIF) of the 16 potent inhibitors and M$^{pro}$.

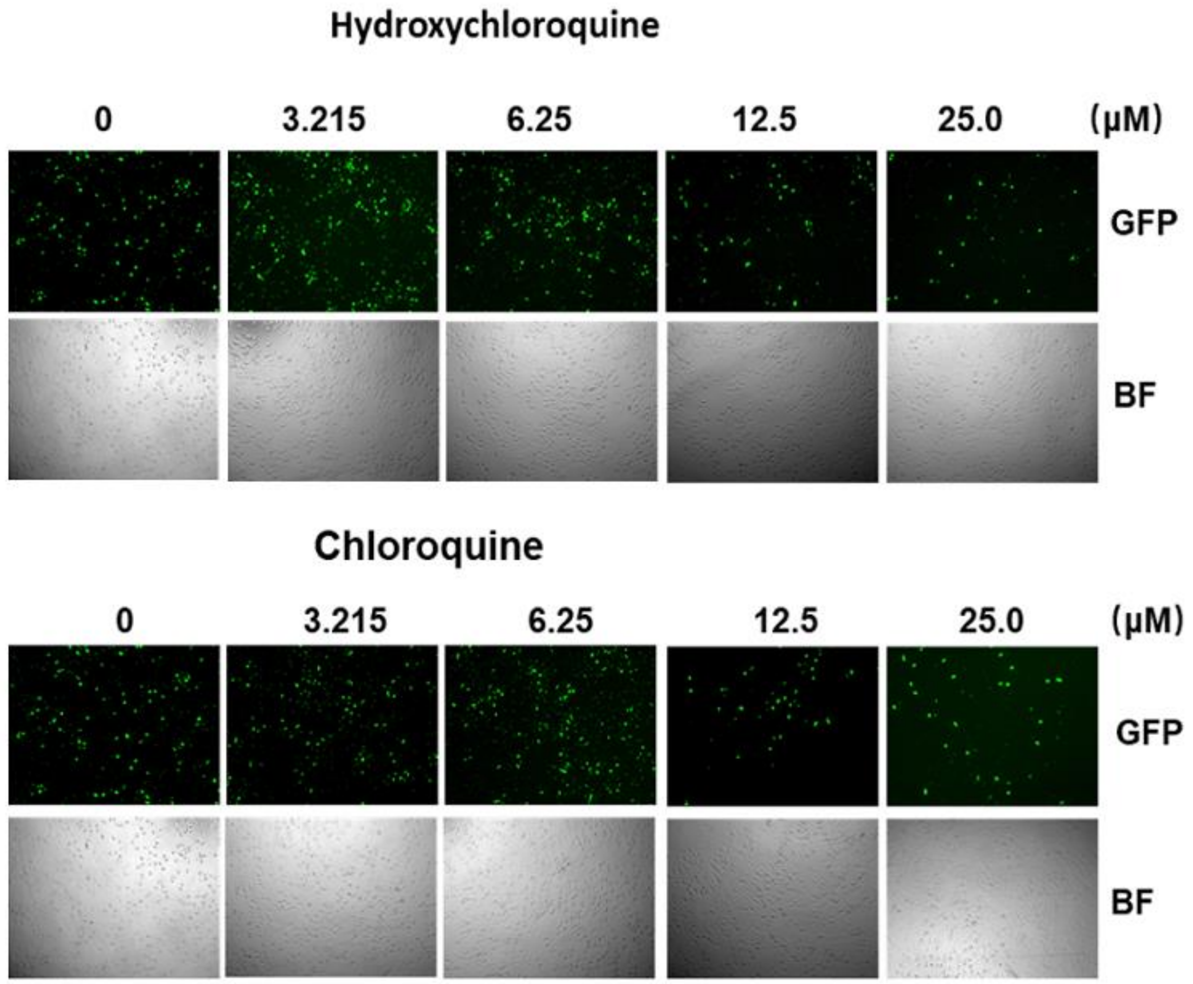


**Figure S10**. Chloroquine and its derivates do not inhibit VSV infection at low concentration. African green monkey kidney cell line VERO was infected with recombinant vesicular stomatitis virus, in which a reporter gene green fluorescence protein (GFP) was inserted into viral genome, at multiplicity of infection (MOI) at 0.01. And then cells were treated with indicated chemicals at different concentration immediately. Ten hours post infection, viral infection was monitored under a microscope. Green fluorescence indicated infected cells. BF, bright field.
